# Transcriptional Signatures of Synaptic Vesicle Genes Define Myotonic Dystrophy Type I Neurodegeneration

**DOI:** 10.1101/2020.07.17.208132

**Authors:** Antonio Jimenez-Marin, Ibai Diez, Garazi Labayru, Andone Sistiaga, Jorge Sepulcre, Adolfo Lopez de Munain, Jesus M. Cortes

## Abstract

Despite significant research, the biological mechanisms underlying the brain degeneration in Myotonic Dystrophy Type I (DM1) remain largely unknown. Here we have assessed brain degeneration by measuring the volume loss (VL) and cognitive deficits (CD) in two cohorts of DM1 patients, and associating them to the large-scale brain transcriptome maps provided by the Allen Human Brain Atlas (AHBA). From a list of preselected hypothesis-driven genes, three of them appear to play a major role in degeneration: dystrophin (*DMD)*, alpha-synuclein (*SNCA)* and the microtubule-associated protein tau (*MAPT)*. Moreover, a purely data-driven strategy identified gene clusters enriched for key biological processes in the central nervous system, such as synaptic vesicle recycling, localization, endocytosis and exocytosis, and the serotonin and dopamine neurotransmitter pathways. Therefore, by combining large-scale transcriptome interactions with brain imaging and cognitive function, we provide a new more comprehensive understanding of DM1 that might help define future therapeutic strategies and research into this condition.

## Introduction

Myotonic Dystrophy type 1 (DM1) is a complex multisystem disease that affects skeletal muscles^1^, the heart^2^, lungs^3^, endocrine system^4^, the regulation of sleep cycles^5^ and other aspects of brain activity^6^. Epidemiologically, DM1 is the most common adult-onset muscular dystrophy in humans, with a reported prevalence of 1/7,400 people worldwide^7^ that is about three times higher in Gipuzkoa^8^, Northern Spain (where this study was performed). Neuroimaging studies has shown the brain damage in DM1 patients, such as gray matter atrophy mainly affecting the frontal and parietal lobes^9^ but also, in the hippocampus^10^ and other subcortical structures^11^. More recent studies, also showed that DM1 produces atrophy along white-matter tracts^12, 13^, affecting the large-scale connectivity of the human brain.

Using an approach that combines magnetic resonance imaging (MRI) and large-scale brain transcriptomics, we aimed here to assess to what extent *the structural damage in the DM1 brain represents a neurogenetic signature.* In contrast to other neurodegenerative diseases in which a large number of candidate genes are implicated, for example about 700 genes in Alzheimer’s Disease (AD)^14^, DM1 is a monogenic disorder caused by a mutation in the gene encoding the myotonic dystrophy protein kinase (*DMPK*)^15^. However, although the disease is monogenic, its phenotype is mainly due to an abnormal activity of the RNA-binding protein muscleblind-like 1 and 2 genes (*MBNL1*, *MBNL2*)^16, 17^ and CUGBP which regulates the expression of many other genes, such as the chloride channel 1 gene (*CLCN1*) that regulates chloride conductance during muscle development^18^, the insulin receptor gene (*INSR*)^19^, the bridging integrator 1 gene (*BIN1*)^20^ or other genes directly related to the main symptoms of the condition^21^. The symptomatology of DM1 is mainly associated to cognitive difficulties including visuospatial processing and executive functioning^22, 23^. Strikingly, the *DMPK* pathogenic genotype has also been associated with other genes that encode proteins implicated in brain neurodegeneration, majorly to tau deposits^24^, but also amyloid beta (Aβ)^25^ or alpha synuclein^26, 27^. Thus, it is suspected that the cognitive deficits and brain damage found in DM1 patients might present some similarities to that in other neurodegenerative diseases, although this issue has yet to be fully addressed. Therefore, despite the monogenic origin of DM1, the gene-to-gene interactome scales up to implicate multiple systems in the brain and body. To date, a precise association between the entire transcriptome and the brain neurodegeneration and cognitive deterioration in DM1 patients remains unexplored.

Some studies have assessed the relationship between genetics and structural brain alterations in DM1, confirming that the number of pathogenic repeats of the cytosine-thymine-guanine (CTG) triplet in the DMPK gene (a parameter used to quantify the molecular severity of the disease) was associated with both stronger gray and white matter atrophy, and to the cognitive deficits of these patients^28, 29^. Here, we performed an intersection analysis of neuroimaging phenotypes and the Allen Human Brain Atlas (AHBA) of large-scale transcriptional human data^30^, following a similar methodology to that used previously^31–33^. Our hypothesis was that identifying the genes whose expression coincided more closely with the brain damage found in DM1 patients, we might better understand the gene relationships associated with the brain damage that arises in this pathology. Similarly, we also assessed the relationship between gene transcription in specific anatomical regions and the brain maps of cognitive deficits in these patients.

## Methods

### Participants, two cohorts

A total of N=95 subjects were recruited to a cross-sectional study, 35 of whom were DM1 patients who were treated at the Neurology Department of the Donostia University Hospital (Gipuzkoa, Spain), while 60 subjects participated as Healthy controls (HCs). All patients and HCs were recruited from the vicinity of the Donostia University Hospital, and the two groups were matched for age, gender and education. The imaging data from the DM1 patients and HCs was acquired at two different Institutions. At one, 19 DM1 patients (mean age 53.3 years [SD ±8.1 years]; 9 males, 10 females) and 29 HC (52.2 [±8.1] years; 12 males, 17 females) were examined and at the second, 16 DM1 patients (48.8 [±7.7] years; 7 males, 9 females) and 31 HC (47.6 [± 7.6] years; 14 males, 17 females). For the mean values, and the comparisons between groups and cohorts see Tables 1 and S1.

**Table 1.** Demographic, clinical and neuropsychological variables. Mean values (standard deviations in brackets) of the different variables separated by cohort. The neuropsychological variables coincide with the different composite-scores from the different cognitive domains. All the neuropsychological variables were calculated using normative data from a healthy Spanish population except for the IQ. Because the average score for each domain is equal to zero, values lower than zero indicate that the mean values in those domains are lower than those in the healthy population (the smaller the value, the worse the performance): bold values indicate p < 0.05. *For gender differences, the Chi^2^ test was used.

|  | *1<sup>st</sup> Cohort* | *2<sup>nd</sup> Cohort* | *t* | *p* | *Effect Size (Cohen)* |
| --- | --- | --- | --- | --- | --- |
| **Demographic** |  |  |  |  |  |
| *N* | 19 | 16 |  |  |  |
| *Age, years* | 53.30 (8.09) | 48.75 (7.72) | 1.69 | .10 | 0.57 |
| *Males, n(%)* | 9 (47.37) | 7 (43.75) | 0.05* | .83 | -0.07 |
| *Education, years* | 16.37 (4.87) | 13.75 (4.85) | 1.59 | .12 | 0.54 |
| **Clinical** |  |  |  |  |  |
| *N* | 19 | 16 |  |  |  |
| *CTG expansion size* | 522.79 (448.44) | 827.13 (433.08) | -2.03 | .05 | -0.69 |
| *MIRS* | 2.47 (0.96) | 3.53 (0.83) | -3.37 | **.002** | -1.17 |
| **Neuropsychological** |  |  |  |  |  |
| *N* | 18 | 16 |  |  |  |
| *Visuospatial* | -0.35 (1.21) | -1.17 (1.34) | 1.86 | .07 | 0.64 |
| *Verbal memory* | -0.04 (2.33) | -0.74 (1.99) | 0.94 | .36 | 0.32 |
| *Attention* | -2.50 (1.92) | -2.62 (2.14) | 0.19 | .85 | 0.06 |
| *Executive functioning* | -0.91 (2.08) | -1.84 (2.40) | 1.21 | .23 | 0.42 |
| *Visual memory* | -0.34 (0.97) | -1.01 (1.14) | 1.83 | .08 | 0.64 |
| *Intelligence (IQ)* | 101.11 (11.51) | 91.19 (13.61) | 2.3 | **.03** | 0.79 |

The DM1 patients were only included if they had molecular confirmation of their DM1 diagnosis, indicating the expansion of hundreds to thousands of CTG repeats in the *DMPK* gene^15^. The diagnosis was obtained when patients were between 18 and 40 years old, and following the adult-onset proposed as the fourth outcome measure for myotonic dystrophy type 1 (OMMYD-4). Patients were excluded if at least one of the following criteria was met: congenital or pediatric disease onset; a history of a major psychiatric or somatic disorder in accordance with DSM-V criteria; acquired brain damage; alcohol or drug abuse; the presence of corporal paramagnetic devices like pacemakers or metal prosthesis that might compromise the MRI studies; and the presence of brain abnormalities that could affect the volumetric analysis. HCs satisfied the same exclusion criteria but the number of the CTG repeats in their *DMPK* gene ranged from 5 to 34^15^. DM1 patients were recruited from the Neuromuscular Unit in the Neurology Department of the Hospital Universitario Donostia, while HCs were recruited from their healthy relatives in whom in the *DMPK* expansion was excluded.

All participants were informed about the study and offered their signed their informed consent. The study was approved by the Ethical Committee of the Donostia University Hospital (code DMRM-2017-01) and it was carried out in accordance with the tenets for human research laid out in the Helsinki Declaration.

### Demographic, clinical and neuropsychological variables

The demographic variables of the subjects recorded were their age, gender and years of education. The clinical variables were the CTG expansion size and Muscular Impairment Rating Scale (MIRS) score^34^. Neuropsychological variables corresponded to composite values from different cognitive domains obtained through a comprehensive neuropsychological evaluation performed by an experienced neuropsychologist who was blind to the patient’s clinical condition (CTG expansion size and MIRS results). The neuropsychological assessment included several subtests from the Wechsler Adult Intelligence Scale III (WAIS III)^35^, including: Digit span, Vocabulary, Block design, Object assembly, Arithmetic, and Similarities. Other cognitive tests used were: Stroop, California Computerized Assessment Package (CALCAP), Raven’s progressive matrices, Rey Auditory Verbal Learning Test (RAVLT)^36^, Word Fluency^37, 38^, Rey-Osterrieth Complex Figure test (ROCF)^39^ and Benton’s Judgement of Line Orientation^40^. The patients’ raw scores were converted into standardized t-values based on the normative scores for the Spanish population in each test. Finally, the different neuropsychological scores were reduced into six different domains: visuospatial (Block design and ROCF copy), verbal memory (RAVLT immediate recall, RAVLT delayed recall, Total RAVLT), attention (Digit span, STROOP word, STROOP color, Simple Reaction Time (RT), election RT, Sequential 1 RT, Sequential 2 RT), executive functioning (Total RAVEN, semantic fluency, phonemic fluency, STROOP color-word, STROOP interference), visual memory (ROCF delayed recall), and intelligence.

### MRI acquisition and preprocessing

For the first cohort, MRI was conducted on a 3 Tesla scanner (TrioTim, Siemens) using a high-resolution 3D sequence of magnetization-prepared rapid acquisition with gradient echo (MPRAGE) and applying the following parameters: Sagittal 3D T1 weighted acquisition, TR = 2300 ms, TE = 2.86 ms, inversion time = 900 ms, flip angle = 9 deg, matrix = 192 x 192 mm^2^, slice thickness = 1.25 mm, voxel dimensions = 1.25 x 1.25 x 1.25 mm^3^, NSA = 1, slices = 144, no gap, total scan duration = 7 min and 22 sec. For the second cohort, MRI was conducted on a 1.5 Tesla scanner (Achieva Nova, Philips), using a high-resolution volumetric turbo field echo (TFE) sequence with the following parameters: Sagittal 3D T1 weighted acquisition, TR = 7.2 ms, TE = 3.3 ms, inversion time = 0 ms, flip angle = 8 deg, matrix = 256 x 232 mm^2^, slice thickness = 1mm, voxel dimensions = 1 x 1 x 1 mm^3^, NSA = 1, slices = 160, no gap, total scan duration = 5 min 34 sec.

To perform voxel-based gray matter (GM) morphometric comparisons between the subjects, DM1 and HCs, we performed Voxel-Based Morphometry (VBM) following a procedure similar to that used previously^41, 42^, an optimized VBM protocol^43^ carried out with the FSL v6.01 software. First, skull-removal was performed, followed by GM segmentation and registration to the MNI 152 standard space using non-linear registration^44^. The resulting images were averaged and flipped along the x-axis to create a left-right symmetric, study-specific GM template. Second, all native GM images were non-linearly registered to this study-specific template and <modulated= to correct for local expansion (or contraction) due to the non-linear component of the spatial transformation. The modulated GM images were then smoothed with an isotropic Gaussian kernel at sigma = 3 and finally, the partial gray matter volume estimates normalized to the subject’s head size were compared.

### Imaging statistical analysis

For the VBM analyses, a generalized lineal model was fitted for each voxel and image using the FSL software, controlling for age and head size with two different contrasts: DM1 < HC and DM1 > HC. All the results were obtained with two-tailed tests, correcting for multiple comparisons using the Monte Carlo simulation clusterwise correction implemented in the AFNI v19.3.00 software, and using 10,000 iterations to estimate the probability of false positive clusters with a p-value < 0.05. Each cohort was analyzed separately and in combination, as explained below, although statistical comparisons between the two cohorts were not performed.

### Transcriptomics brain maps

To build brain maps of transcription we took advantage of the publicly available data from the Allen Human Brain Atlas (AHBA – http://human.brain-map.org/)^30^. The dataset consisted of MRI images and a total of 58,692 microarray-based transcription profiles of about 20,945 genes sampled over 3,702 different regions across the brains of six humans. To pool all the transcription data into a single brain template, we followed a similar procedure to that employed elsewhere^45^. First, to re-annotate the probes to genes we made use of the re-annotator toolkit^46^.

Second, we removed those probes with insufficient signal by looking at the sampling proportion (SP), which was calculated for each brain as the ratio between the samples with a signal greater than the background noise divided by the total number of samples. Probes with a SP lower than 70% in any of the six brains were removed from the analysis, thereby ensuring sufficient sampling power in all the brains. After that, we chose the value of the probe for each gene with the maximum differential stability (DS), accounting for the reproducibility of gene expression across brain regions and individuals, and calculated using spatial correlations similar to those employed previously^47^. For this the Automated Anatomical Labelling (AAL) atlas was used^48^ from which the cerebellum was excluded, resulting in 90 different anatomical regions*. Finally, to remove the inter-subject differences the transcription values for each gene and brain were transformed into Z-scores, and pooled together from the six different brains, obtaining a single map using the MNI coordinates provided in the dataset. Finally, to eliminate the spatial dependencies of the transcription values at the sampling sites (that is, to correct for the fact that nearest sites have more correlated transcription), we finally obtained a single transcription value for each region in the AAL atlas by calculating the median of all the values belonging to the given region.

### Association between volume loss and transcriptomics

Volume loss (VL) was defined as the t-statistic resulting from the group comparison DM1 < HC. To associate VL with transcriptomics, we transformed the t-statistics map to the same AAL atlas as that used for the transcription values, calculating the median of the t-statistics between all values in each region. For each gene, we calculated a similarity index using the Pearson correlation coefficient between VL and transcriptomics, the two variables represented in vectors with a dimension equal to the number of regions in the atlas. This procedure was done separately for the two cohorts.

### Association between cognitive deficits and transcriptomics

For the association between cognitive deficits (CD) and transcriptomics, we first built brain maps of CD using the BrainMap meta-analysis platform (http://www.brainmap.org/). In particular, the Sleuth tool v3.0.3^49^ was used to search all the papers in the database using the name of each neuropsychological domain as a keyword. In this way, all the co-activation coordinates that resulted from studies based on functional imaging when the participant in the scanner was performing a task related to each neuropsychological domain were obtained. Next, the GingerAle tool v3.0.2^50^ was used to pool all the coordinates onto a single co-activation brain-map for each neuropsychological domain, representing these as a Z-score after applying the activation likelihood estimation (ALE) method. The brain maps were then transformed to the same AAL atlas by calculating the median of all the Z-scores belonging to the same region in the atlas. Finally, the association between CD and transcriptomics was assessed for each gene through the similarity index, equivalent to the Pearson correlation coefficient between CD and transcriptomics, the two variables were represented in the vectors with a dimension equal to the number of regions in the atlas. Transcriptomics correlates were only obtained for those neuropsychological domains that were most affected, identified by choosing a mean Z-score < −2, previously standardized to the normative scores based on a Spanish population. This procedure was done separately for the two cohorts.

### DM1 relevant genes and gene ontology

Relevant genes were identified by combining the results from two different strategies. The first was the hypothesis-driven strategy (HDS) that involved reviewing previous studies to identify genes that play a relevant role in some neurobiological aspects of DM1 (Table S2). The second was the data-driven strategy (DDS), which consists of identifying the genes with the strongest transcription correlation with the parameter of interest, either VL or CD. In particular, we chose those genes with similarity index values where *z* < −2 or *z* > 2, i.e.: outliers of the correlation distribution in both the negative and positive tails. Genes in the positive tail (*z* > 2) were designated as *pos-corr* genes (P), whereas those in the negative tail (*z* < −2) were considered *neg-corr* genes (N). For example, when assessing the VL, the P-genes were systematically expressed more strongly than other genes in the brain regions with more pronounced atrophy, whereas the N-genes were expressed much more weakly than the rest. Although this procedure was performed separately for the two cohorts, the P- and N-genes finally adopted were those present in the two cohorts. We also considered those genes that were expressed similarly to the P- and N-genes as relevant genes, which were dubbed *connector hubs* (C). To identify these, we made use of the “gene-expression connectivity matrix” that represents the similarity in expression between gene pairs obtained from the Pearson pairwise correlations for each entry^†^. For those genes absent in the groups of P- and N-genes, the strength towards the two P- and N-tails was calculated separately. PC-genes were identified as those genes with a Z-score in connectivity strength towards the P-tail >2 and similarly, NC-genes were identified by having a Z-score in the strength towards the N-tail > 2. For the hypothesis-driven genes given in Table S2 that were defined as P or N genes, their statistical significance was assessed by surrogate-data testing. The BrainSMASH tool^51^ was used to build null-distributions by generating 10,000 random maps with the same spatial autocorrelation as that for VL or CD.

Finally, we pooled together all the relevant genes (P-genes, N-genes, PC-genes, NC-genes and the genes that were common to PC and NC) to perform unsupervised K-means clustering with the *Silhouette* strategy to identify the optimal number of clusters. For each cluster, we performed a gene ontology *GO biological process*^52^ and *Reactome pathways*^53^ overrepresentation test using PANTHER v15.0 (http://pantherdb.org/), with the entire Homo Sapiens genome as the reference list and applying a Fisher’s Exact test with Bonferroni correction (p < 0.05). To make the data more readily interpretable, we only reported ≥2 fold enrichment.

## Results

A total of 35 DM1 patients and 60 HCs participated in this study, examined in two different cohorts at two distinct centers. In the first of these, 19 DM1 patients were recruited (9 males) with an average age of 53.3 years (range 42.1 – 69.5 years) and with 16.4 years of education (range 8 – 25 years). With respect to the clinical variables, these patients had an average MIRS score of 2.5 (range 1 – 4) and a mean CTG expansion of 522.8 (range 65 – 1733). These patients were compared with 29 well-matched HCs. In the second cohort, 16 DM1 patients were recruited (7 males), with an average age of 48.8 years (range 36.0 – 61.0 years) and 13.2 years of education (range 6 – 22 years). The mean MIRS score in this cohort was 3.5 (range 2 – 5) and with a CTG expansion size of 827.1 (range 267 – 1833). These patients were compared to 31 well-matched HCs, and all the demographic, clinical and neuropsychological variables collected in the study are detailed in Tables 1 and S1. As explained in the Methods, the two cohorts were considered independently for the imaging analyses, obtaining the corresponding VL brain maps and identifying the list of relevant genes for each cohort separately. Subsequently, the two lists of genes were assessed and only the genes common to the two cohorts were analyzed.

The VL brain maps from each cohort consisted of several widespread cortical and subcortical regions (Figure 1A). There was bilateral subcortical VL in the hippocampus, thalamus, basal ganglia and cerebellum, while cortical VL was evident in multiple regions that included part of the temporal, occipital, parietal cortices, precuneus, pars orbitalis and medial prefrontal cortex (for details see Table S3 and S4). Importantly, the VL maps for the two cohorts differed more in the cortical regions (Dice index, D = 0.3) and they were more similar in the subcortical ones (D = 0.8), indicating that subcortical atrophy was more prevalent in DM1 and less variable than cortical atrophy.

**Figure 1.**
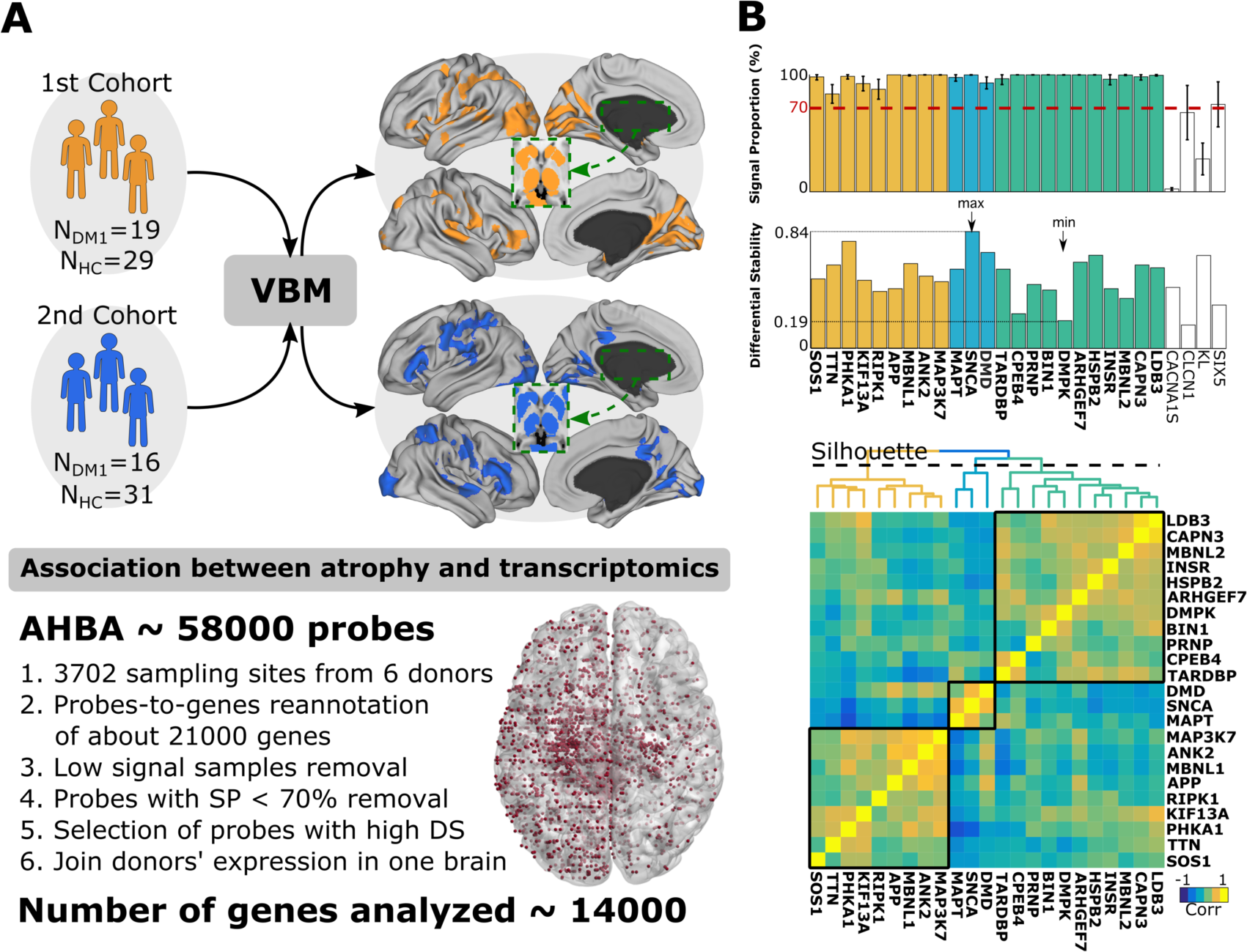
Methodological scheme for the association between transcriptomics and atrophy in DM1, measured as brain volume loss (VL). **A:** Two cohorts of DM1 patients were recruited (orange and blue) and we obtained the brain maps of the VL for each, comparing the images with a group of HCs using Voxel-Based-Morphometry (VBM), correcting for multiple comparisons. We aimed to characterize the association between VL and transcriptomics, assessing the similarity in the spatial patterns of VL across brain regions and the spatial patterns of gene transcription from the AHBA dataset, preprocessed following a pipeline that is summarized in six main steps (for further details, see methods). After running the AHBA pipeline, about 14K genes finally had transcription values used in the analysis from the 58K probes originally available. The red regions in the brain correspond to the sites at which transcription was sampled. **B:** The sampling proportion (SP) and differential stability (DS) for the 27 preselected hypothesis-driven genes included in the list of candidates relevant to DM1 (obtained by reviewing the literature). The *CACNA1S, CLCN1, KL* and *SIX5* genes did not have a mean SP value above 70% and thus, they were excluded from further analyses. Of the remaining 23 genes, the maximum DS corresponded to *SNCA* and the minimum DS to *DMPK* (see arrows). By examining the spatial similarity in the transcription values, the remaining 23 candidate genes were clustered into three groups. The blue one formed by *DMD*, *SNCA* and *MAPT* played a major role in the characterization of VL. Abbreviations: Muscular Dystrophy Type I (DM1); Healthy Control (HC); Voxel-based morphometry (VBM); Allen Human Brain Atlas (AHBA); Sampling proportion (SP); Differential Stability (DS).

The main steps in the pipeline were followed to analyze the transcriptomics through AHBA dataset (Figure 1A), providing brain maps of transcriptional activity for each gene that were spatially-correlated with the brain maps of VL and subsequently, with those of CD. A list of the 27 most relevant hypothesis-driven genes in DM1 was drawn up (Figure 1B, for details on references supporting the selection of each gene see Table S2). Of these 27 preselected genes, 4 genes were discarded based on their SP and DS values (*KL*, *SIX5*, *CLCN1*, *CACNA1S)*. These 4 genes had less than 70% SP (Figure 1B), which implied insufficient transcription signal across all the sampling sites. Therefore, the final list of the most relevant hypothesis-driven genes contained: *DMPK*, *HSPB2*, *INSR*, *CPEB4, ANK2, ARHGEF7, SOS1, PHKA1, MBNL1, KIF13A, APP, MAPT, SNCA, MBNL2, RIPK1, PRNP, TARDBP, MAP3K7, BIN1, DMD, LDB3, TTN* and *CAPN3.* When the pairwise gene-to-gene similarity in transcription was evaluated across the brain for these 23 genes, Silhoutte maximization identified three clusters (in yellow, blue and green in Figure 1B), indicative of functional similarities in the transcription signals among the 23 most-relevant genes.

We next applied a DDS that involved identifying the genes from the entire transcriptome with maximal association in the VL maps, resulting in a total of 370 N-genes and 187 P-genes from the 1st cohort, and 441 N-genes and 161 NP-genes from the 2nd one. The genes common to the two cohorts were those finally used in the analysis, a total of 251 N-genes and 101 P-genes (Figure 2A). Interestingly, two of the genes in the list of the 23 most relevant hypothesis-driven genes also appeared in the list of N-genes, *SNCA* and *DMD*, displaying a similar transcription pattern as the *MAPT* gene (blue cluster in Figure 1B). The spatial-correlation of *SNCA* and *DMD* transcription with the amount of VL in the different brain regions proved to be negative for the two genes and smaller in the two cohorts: < −0.52 for *DMD* and < −0.70 for *SNCA* (Figure 2B and Table S5). In addition to identifying the N- and P-genes, we also searched for the NC- and PC-genes (Figure 3A) that represent *hubs* towards the N and P tails of the expression similarity matrix. We found 452 NC-genes, 396 PC-genes, and 238 genes connecting both N- and P-tails. Remarkably, in the group of PC genes we found *LDB3*, *CAPN3* and *HSPB2* that were in the list of hypothesis-driven genes, belonging to the same cluster of expression similarity (green cluster, Figure 1B). In addition, the preselected gene *PHKA1* was also common to the groups of NC and PC genes.

**Figure 2.**
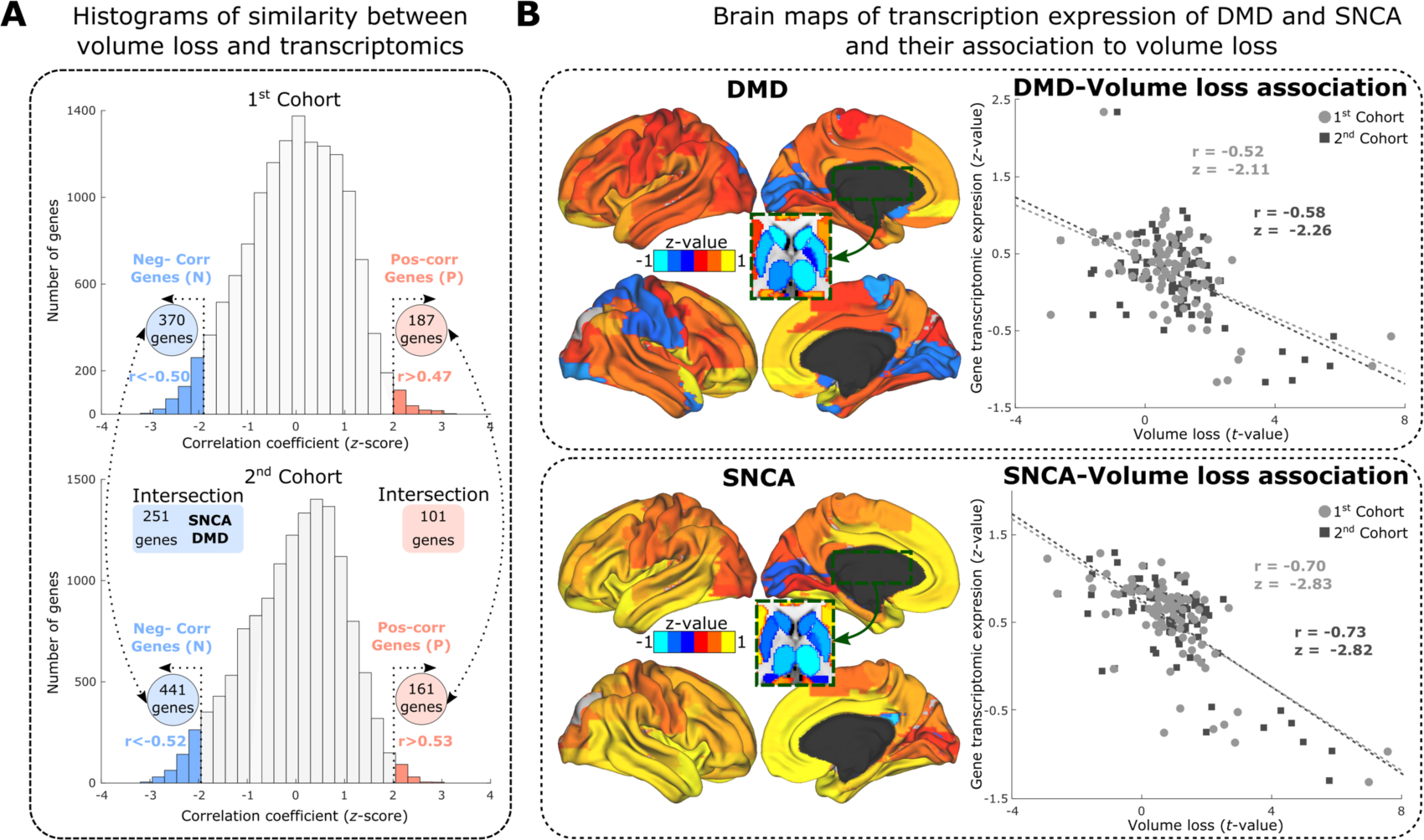
Data-driven strategy to determine the association between the transcriptome and VL in DM1. **A:** Histogram of the spatial-correlation values (measured as the Z-score) between volume loss (VL) and transcriptional activity for all the genes in both cohorts. For both cohorts the N-genes (*z* < −2) and P-genes (*z* > 2) are colored in blue and red, respectively. The final list of genes used for further analyses are those that are common to the two cohorts, consisting of 251 N-genes and 101 P-genes. From all the genes that provide a maximum association between VL and transcriptomics (Table S6 and S7), only two genes were in the panel of preselected genes: *SNCA* and *DMD*. **B:** Brain maps of transcription in the brain regions for the two genes *DMD* and *SNCA*, which provided a high spatial correlation (r) with the VL brain maps for both cohorts.

**Figure 3.**
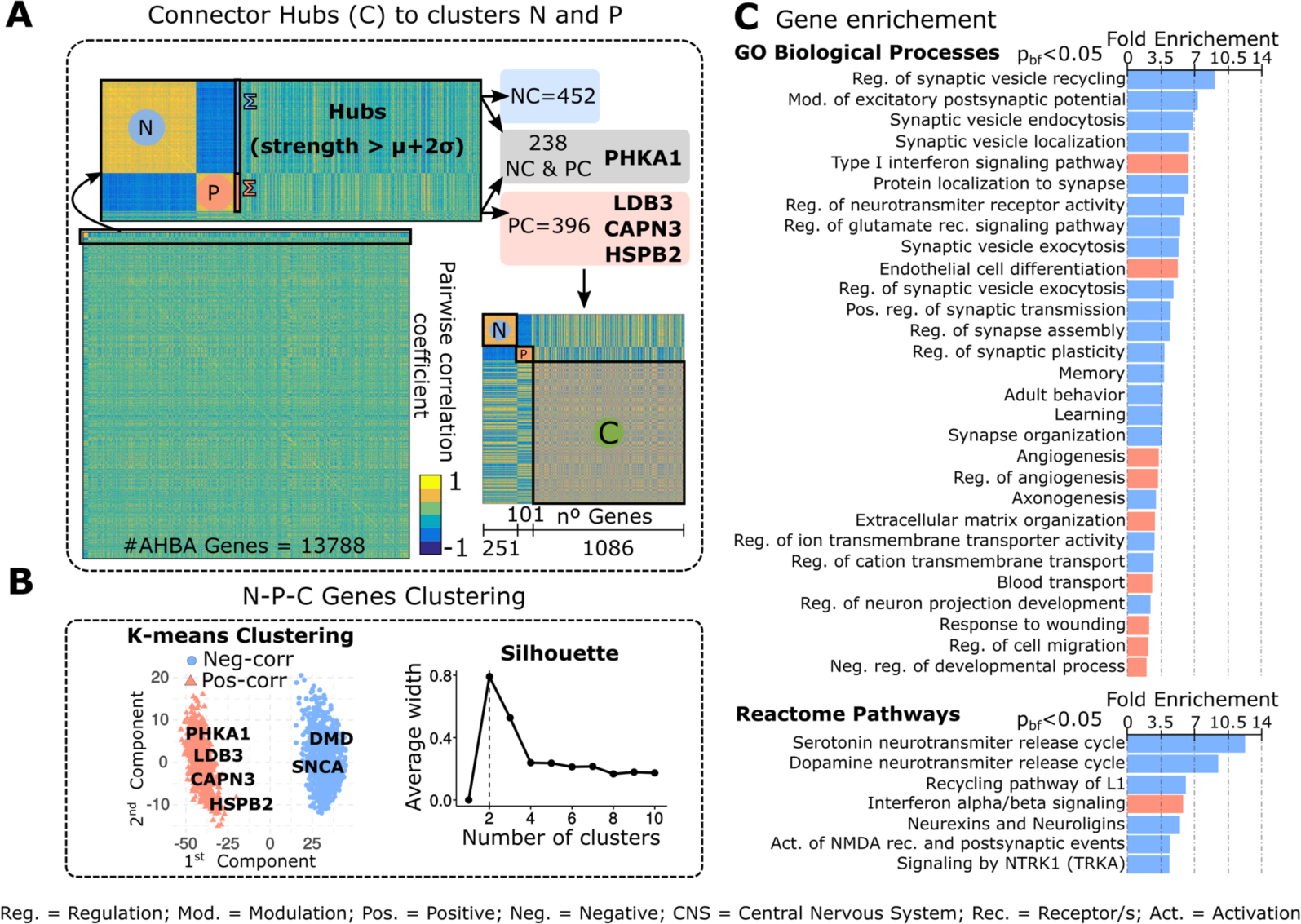
Functional description of the genes with the highest association with volume loss (VL). **A:** An all-to-all gene-expression similarity matrix identified the connector hub genes. A total of 1086 genes were found, equal to the sum of the NC = 452 (blue), PC = 396 (red) and 238 common NC and PC genes (gray). **B:** Two clusters were finally found that pooled all gene classes, the blue one contains the original N=251 genes and the red one containing the original P=101 genes. **C:** Gene enrichment for *GO biological process* and *Reactome pathways:* in blue are the neg-corr genes and in red, the pos-corr genes. Abbreviations: Regulation (Reg.); Modulation (Mod.); Positive (Pos.); Negative (Neg.); Central Nervous System (CNS); Receptor/s (Rec.); Activation (Act.).

Pooling the N, P, NC and PC genes together, along with those common to both the NC and PC categories, we adopted a DDS to achieve unsupervised clustering of the expression similarity matrix, identifying two major clusters after Silhouette maximization (in blue and red in Figure 3B). Importantly, the N and P genes fully segregated into the two differentiated clusters, with all the N-genes belonging to the blue cluster and all the P-genes to the red one, thereby confirming the different functional roles of the groups of genes in the N- and P-tails. These two clusters were used separately for gene enrichment analysis (the list of genes included in each cluster are given in Table S6 and S7). The search for the *GO biological processes* and *Reactome pathways* confirmed the differentiated roles of these two clusters, with the *neg-corr* genes more related to neuronal and synaptic function, involving key synaptic vesicle events such as recycling, localization, endocytosis and exocytosis but also, the dynamics of serotonin and dopamine neurotransmitter release (Figure 3C). By contrast, the cluster of pos-corr genes was more related to non-neuronal activities, such as interferon signaling, endothelial cell differentiation, angiogenesis, blood transport and cell development.

To assess how the transcriptomics correlated with CD, we focused on the neuropsychological domains in which the composite score reflected strong impairment, satisfying *z* < −2, which was only the case for the attention category (1^st^ cohort *z* = −2.5, 2^nd^ cohort *z* = −2.62: see Table 1 and Figure 4A for the distribution of all the Z-score values from the two cohorts). As indicated in the Methods, the attention composite was obtained by averaging the Z-scores for the following domains: Digit span, STROOP word, STROOP color, Simple RT, election RT, Sequential 1 RT, Sequential 2 RT.

**Figure 4.**
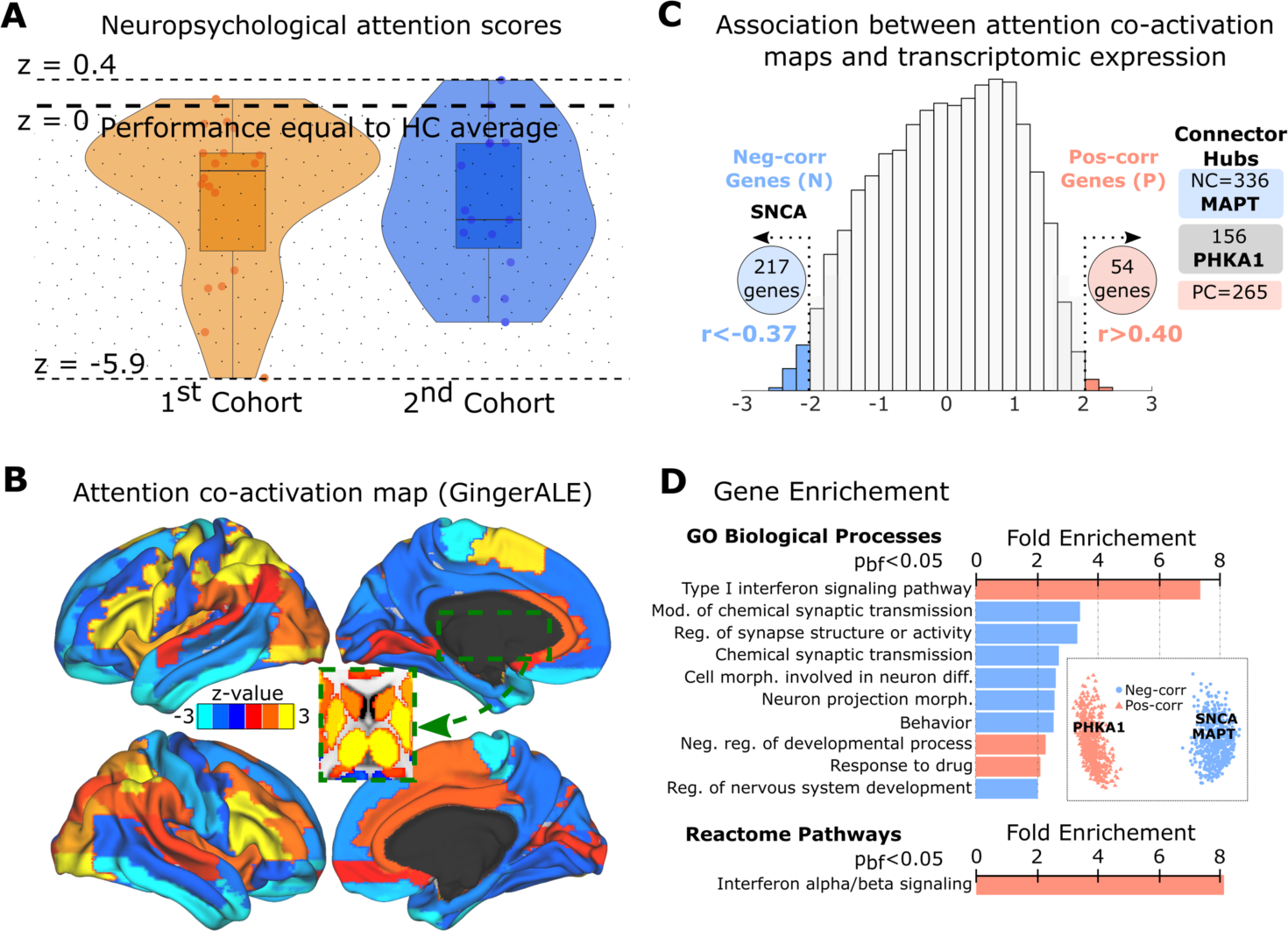
Data-driven strategy to define the association between the transcriptome and attention co-activation maps in DM1. **A:** Attention scores measured as Z-scores for the two cohorts. Because the Z-scores were normalized to the values in the HCs, negative values of *z* indicate worse performance than the HCs. **B:** Attention co-activation maps built with the GingerALE tool and projected onto the atlas. **C:** Histograms of the spatial correlations between the Z-scores of the attention maps and the transcriptional activity for each gene. The tail of N-genes (*z* < −2, colored in blue) includes the *SNCA* gene from the list of preselected genes, whereas the tail of the P-genes (*z* > 2, red) does not include any of these. Following a procedure similar to that described in Figure 3A, we identified the PC-genes (red), NC-genes (blue, including *MAPT*), and those common to the NC and PC (gray, including *PHKA1*). **D:** After pooling all classes of genes together and clustering, two groups were defined: one including all the neg-corr genes (blue, with *SNCA* and *MAPT)* and one with the pos-corr genes (red, with *PHKA1)*. Gene enrichment for the tags *GO biological process* and *Reactome pathways*. As in Figure 3C, the two clusters also represented two separated functions: the neg-corr one correlated with neuronal functions, whilst the pos-corr correlated to non-brain functions. Abbreviations: Regulation (Reg.); Modulation (Mod.); Positive (Pos.); Negative (Neg.).

We then obtained brain co-activation maps using “Attention” as a keyword for the search in GingerAle (Figure 4B). As for VL, we calculated the association between the attention co-activation maps and transcriptomics by calculating the spatial Pearson correlation between the two Z-score vectors (one value per brain region in the atlas, see the histogram of all the correlations in Figure 4C). There were a total of 217 N-genes (including *SNCA*), 54 P-genes, 336 NC-genes (including *MAPT* from the list of preselected hypothesis-driven genes), 265 PC-genes, and 156 genes common to both the NC- and PC-gene sets (including *PHKA1)*. Pooling together all classes of genes, we followed a DDS of unsupervised clustering and identified two clusters after Silhoutte maximization (Figure 4C, Tables S8 and S9). Like VL, the two clusters were highly segregated and incorporated all *neg-corr* genes in one cluster, which was enriched in genes related to neuronal and synaptic function (Figure 4D), with all the *pos-corr* genes in the other cluster enriched in non-brain related activities.

## Discussion

The AHBA provides information on the transcriptome across the brain in unprecedented detail, covering about 3,702 sampling sites and allowing activity patterns to be built for about 20,500 genes as a specific signature for each anatomic region. To date, its use has shed some light on several fundamental aspects of the brain and their association with the transcriptome, such as myelination^54^, hierarchical cortical organization^55^, visuomotor integration^33^ or large-scale connectivity^31, 56, 57^. In addition it has provided data regarding pathologies, suggesting novel molecular mechanisms underlying some disorders, such as Autism Spectrum Disorder^58^ or functional neurological disorder^59^. It is important to emphasize that our methodology, assessing the relationships between brain images associated with neurodegeneration and the entire transcriptome is complementary to other techniques, such as genome-wide association studies (GWAS)^60^, simultaneously addressing genotype–phenotype associations from hundreds of thousands to millions of genetic variants in a data-driven manner. However, it is also important to note that such a vast number of multiple-comparisons requires very large samples to achieve statistical power, which is an important limitation for monogenic disorders like DM1.

Monogenic disorders are paradigmatic disease-models of neurogenetic origin, whereby a mutation in a single gene can cause the disease. However, the alterations causing DM1 appear to propagate to a larger proportion of the gene-to-gene interactome, affecting the function of several other genes. Here, we adopted a novel approach using the AHBA to study the transcriptomic correlates of brain damage and CD in DM1 patients, which to our knowledge has yet to be addressed. Our first finding is related precisely to the mutation located in the untranslated region of the DMPK gene, which triggers the disease and that we found to have the lowest DS (0.19) among the panel of the 23 hypothesis-driven preselected genes. Thus, *DMPK* expression was less reproducible in different brain regions and individuals, and consequently, it was more poorly enriched in terms of brain-related biological processes^47^. By contrast, the *SNCA* gene has the highest DS value (0.84). Our second finding is related to the clusters of spatial similarity among the hypothesis-driven genes in DM1. In particular, we found three clusters, one including the *DMPK* gene together with *LDB3*, *CAPN3* and *HSPB2*, these three latter genes associated with VL in DM1 patients. The second cluster contains several genes with a similar expression to *PHKA1,* which was associated with VL and CD in these patients. Finally, the third cluster contains the genes *SNCA*, *DMD* and *MAPT*, and as we show, this cluster plays the most relevant role in VL and CD in DM1.

A DDS revealed the existence of two tails in the distribution of spatial-similarity between transcription activity and VL or CD. Importantly, the three *SNCA*, *DMD*, *MAPT* genes were present in the negative tail (*z* < −2) of neg-corr genes, indicating that in those brain regions where more VL or CD existed the three genes are systematically transcribed more weakly, and vice versa. Thus, in the regions where the three genes are more strongly expressed, VL or CD was less pronounced. Moreover, this cluster was enriched towards brain-related biological activities, such as synaptic vesicle dynamics (recycling, localization, endocytosis and exocytosis), and serotonin and dopamine neurotransmitter release. By contrast, the pos-corr genes belong to the positive tails (*z* > −2) and they were enriched towards non-brain related functions, mainly interferon signaling, endothelial cell differentiation, angiogenesis or blood transport, processes known to be affected in DM1^2, 61–63^.

The DDS also revealed novel relationships between the *MAPT*, *SNCA* and *DMD* genes themselves, providing instructions to produce the proteins tau, alpha-synuclein and dystrophin, respectively. These predictions are consistent with previous studies on DM1 patients, for instance the detection of alpha-synuclein Lewy bodies^27^, splicing abnormalities in dystrophin^64, 65^ and tau-positive degenerative neurites (but not Aβ plaques)^66^, and more recently it was suggested the molecular pathways that associate the two *DMPK* and *MAPT* genes^24^ may be involved in this condition.

The DDS also revealed new genes implicated in DM1, initially absent from the hypothesis-driven list and enriched in brain-related functions, mainly affecting synaptic vesicles or the dopamine and serotonin pathways (Table S10). From the full list of statistically significant contributions, it is important to note that the findings connecting DM1 with key biological processes in the CNS are in full agreement with previous studies showing synaptic protein dysregulation^67, 68^, events mediated by *RAB3A* upregulation and *SYN1* hyperphosphorylation in transgenic DM1 mice, and also in transfected cells and post-mortem brains of DM1 patients. Moreover, alterations to synaptic proteins have also been seen to cause behavioral and electrophysiological dysfunction, affecting neurotransmitter signaling, and reducing the dopamine and 5-hydroxyindoleacetic acid (a serotonin metabolite) availability. These data are consistent with our results that DM1 is related to alterations in short-term synaptic plasticity and neurochemical functioning.

In conclusion, we have studied two different cohorts of DM1 patients, each one well-matched to a group HCs, and by employing a DDS that addresses all the hypothesis driven preselected genes for DM1, *DMD*, *SNCA* and *MAPT* were seen to have a major influence on brain damage and CD. Moreover, we found an enrichment of key biological processes in the CNS, such as synaptic vesicle cycling, recycling and dynamics, and also serotonin and dopamine neurotransmitter signaling. Further studies should clarify whether the interactions between *DMD*, *SNCA* and *MAPT* can be generalized to other degenerative or developmental conditions, or if these are specific to DM1.

## Acknowledgments

We wish to thank Prof. Virginia Arechavala for providing us with an updated list of relevant genes in DM1, some of which were considered in our study. J.M.C is funded by Ikerbasque: The Basque Foundation for Science and from the Ministerio de Economia, Industria y Competitividad (Spain) and FEDER (grant DPI2016-79874-R), and from the Department of Economic and Infrastructure Development of the Basque Country (Elkartek Program, KK-2018/00032 and KK-2018/00090). A.L.d.M was founded by the Institute of Health Carlos III co-founded by Fondo Europeo de Desarrollo Regional-FEDER (grant PI17/01841), CIBERNED (grant 609), and La Caixa Foundation (grant HR17-00268). A.S was founded by the Institute of Health Carlos III co-founded by Fondo Europeo de Desarrollo Regional-FEDER (grant PI17/01231), and the Basque Government (grant SAIO08-PE08BF01). A.J.M was partially funded by Euskampus Fundazioa and a predoctoral grant from the Basque Government (PRE_2019_1_0070). G.L was founded by a predoctoral grant from the Basque Government (PRE_2016_1_0187).

## Supplementary Information

### Comparison of the two cohorts

When comparing the demographic, clinical and neuropsychological variables between the two cohorts, we found significant differences in MIRS (p=0.002) and IQ (p=0.03), showing respectively higher MIRS and lower IQ in the second cohort as compared to the first. The differences in CTG expansion size and age were almost significant, with effect size equal to −0.69 and 0.56 respectively, indicating a tendency of higher CTG expansion size and lower age in the second cohort as compared to the first. When comparing the neuropsychological scores between cohorts, no significance differences were found in any of the cognitive domains. Notice that, all the domains had a z-score lower than zero, meaning that the patients in the two cohorts had worse performance than the healthy controls. The more affected domain was the attention, that had a z-score lower than −2 in the two cohorts.

**Table S1.** **Age, gender and years of education between DM1 patients and HCs.** Mean values of different variables are shown (standard deviations are given in brackets). *For gender differences, the Chi2 test was used.

|  | *DM1* | *HC* | *t* | *p* | *Effect Size (Cohen)* |
| --- | --- | --- | --- | --- | --- |
| *1st Cohort* |  |  |  |  |  |
| *N* | 19 | 29 |  |  |  |
| *Age, years* | 53.30 (8.09) | 52.17 (8.05) | 0.47 | .64 | 0.14 |
| *Males, n(%)* | 9 (47.37) | 12 (41.38) | 0.17* | .68 | -0.12 |
| *Education, years* | 16.37 (4.87) | 14.79 (3.36) | 1.33 | 0.19 | 0.39 |
| *2nd Cohort* |  |  |  |  |  |
| *N* | 16 | 31 |  |  |  |
| *Age, years* | 48.75 (7.72) | 47.55 (7.54) | 0.51 | .61 | 0.16 |
| *Males, n(%)* | 7 (43.75) | 14 (45.16) | 0.01* | .93 | -0.03 |
| *Education (years)* | 13.75 (4.85) | - | - | - | - |

**Table S2.** **List of relevant genes for the neurobiological aspects of DM1 obtained after reviewing previous studies.** Red rows indicate that these genes were not included in the transcriptomics analyses because they had SP < 70%, in at least one of the donors’ brain.

| *HGNC Symbol* | *Human Symbol* | *References* |
| --- | --- | --- |
| *DMPK* | DM1 protein kinase | (1–5) |
| *HSPB2* | DMPK-Binding Protein | (6) |
| *INSR* | Insulin receptor | (7) |
| *CPEB4* | Cytoplasmic polyadenylation element binding protein 4 | (8) |
| *ANK2* | Ankyrin 2 | (9) |
| *ARHGEF7* | Rho guanine nucleotide exchange factor 7 | (10) |
| *SOS1* | SOS Ras/Rac guanine nucleotide exchange factor 1 | (11) |
| *PHKA1* | Phosphorylase kinase regulatory subunit alpha 1 | (12) |
| *MBNL1* | Muscleblind-like splicing regulator 1 | (13, 14) |
| *KIF13A* | Kinesin family member 13A | (11, 15) |
| *APP* | Amyloid beta precursor protein | (16) |
| *MAPT* | Microtubule associated protein tau | (17) |
| *SNCA* | Synuclein alpha | (18, 19) |
| *MBNL2* | Muscleblind like splicing regulator 2 | (20, 21) |
| *RIPK1* | Receptor interacting serine/threonine kinase 1 | (22) |
| *KL* | Klotho | (23, 24) |
| *PRNP* | Prion protein | (25) |
| *TARDBP* | TAR DNA binding protein | (26, 27) |
| *SIX5* | SIX homeobox 5 | (28, 29) |
| *MAP3K7* | Mitogen-activated protein kinase kinase kinase 7 | (30) |
| *BIN1* | Bridging integrator 1 | (31) |
| *CLCN1* | Chloride voltage-gated channel 1 | (32) |
| *DMD* | dystrophin | (33–37) |
| *LDB3* | LIM domain binding 3 | (38) |
| *TTN* | Titin or connectin | (39, 40) |
| *CACNA1S* | calcium voltage-gated channel subunit alpha1 A | (41) |
| *CAPN3* | Calpain 3 | (39) |

**Table S3.**
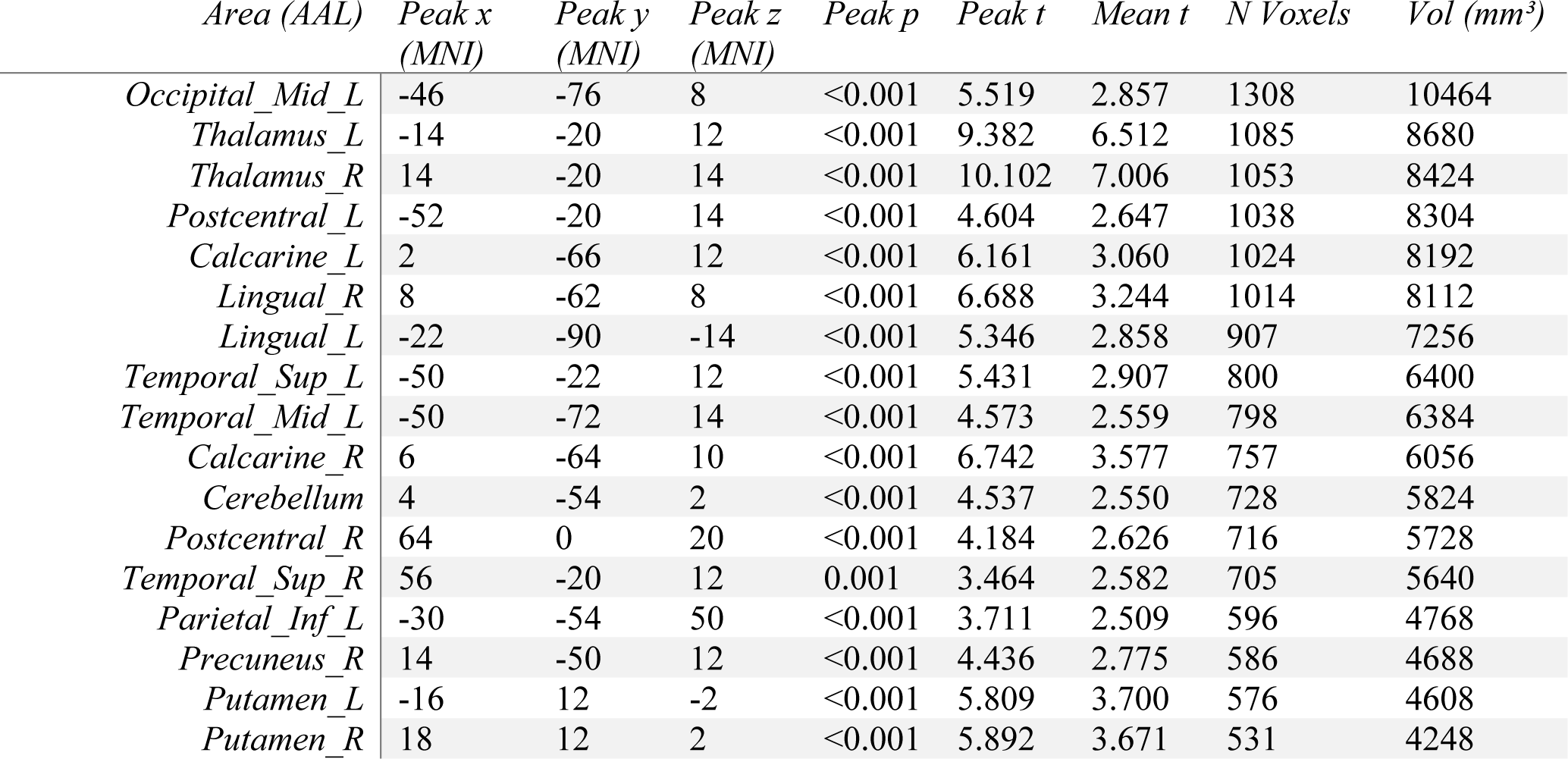
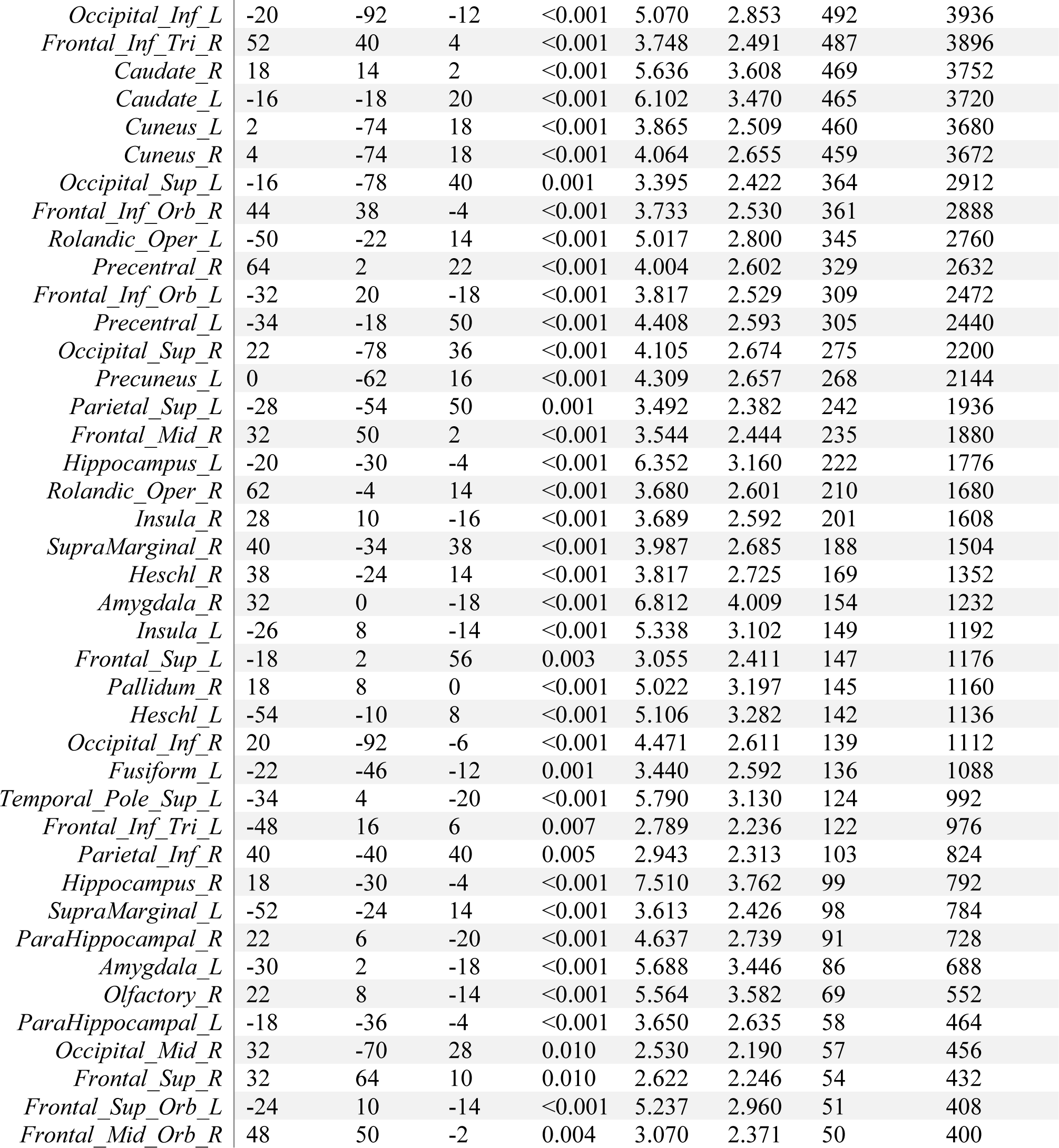
**Anatomical characterization of atrophied regions in the 1st Cohort.** We only represented those regions having more than 50 voxels of overlapping between with the map of volume loss, resulting from the group comparison contrast DM1 < HC.

| *Area (AAL)* | *Peak x*<br>*(MNI)* | *Peak y*<br>*(MNI)* | *Peak z*<br>*(MNI)* | *Peak p* | *Peak t* | *Mean t* | *N Voxels* | *Vol (mm<sup>3</sup>)* |
| --- | --- | --- | --- | --- | --- | --- | --- | --- |
| *Occipital_Mid_L* | -46 | -76 | 8 | <0.001 | 5.519 | 2.857 | 1308 | 10464 |
| *Thalamus_L* | -14 | -20 | 12 | <0.001 | 9.382 | 6.512 | 1085 | 8680 |
| *Thalamus_R* | 14 | -20 | 14 | <0.001 | 10.102 | 7.006 | 1053 | 8424 |
| *Postcentral_L* | -52 | -20 | 14 | <0.001 | 4.604 | 2.647 | 1038 | 8304 |
| *Calcarine_L* | 2 | -66 | 12 | <0.001 | 6.161 | 3.060 | 1024 | 8192 |
| *Lingual_R* | 8 | -62 | 8 | <0.001 | 6.688 | 3.244 | 1014 | 8112 |
| *Lingual_L* | -22 | -90 | -14 | <0.001 | 5.346 | 2.858 | 907 | 7256 |
| *Temporal_Sup_L* | -50 | -22 | 12 | <0.001 | 5.431 | 2.907 | 800 | 6400 |
| *Temporal_Mid_L* | -50 | -72 | 14 | <0.001 | 4.573 | 2.559 | 798 | 6384 |
| *Calcarine_R* | 6 | -64 | 10 | <0.001 | 6.742 | 3.577 | 757 | 6056 |
| *Cerebellum* | 4 | -54 | 2 | <0.001 | 4.537 | 2.550 | 728 | 5824 |
| *Postcentral_R* | 64 | 0 | 20 | <0.001 | 4.184 | 2.626 | 716 | 5728 |
| *Temporal_Sup_R* | 56 | -20 | 12 | 0.001 | 3.464 | 2.582 | 705 | 5640 |
| *Parietal_Inf_L* | -30 | -54 | 50 | <0.001 | 3.711 | 2.509 | 596 | 4768 |
| *Precuneus_R* | 14 | -50 | 12 | <0.001 | 4.436 | 2.775 | 586 | 4688 |
| *Putamen_L* | -16 | 12 | -2 | <0.001 | 5.809 | 3.700 | 576 | 4608 |
| *Putamen_R* | 18 | 12 | 2 | <0.001 | 5.892 | 3.671 | 531 | 4248 |
| *Occipital_Inf_L* | -20 | -92 | -12 | <0.001 | 5.070 | 2.853 | 492 | 3936 |
| *Frontal_Inf_Tri_R* | 52 | 40 | 4 | <0.001 | 3.748 | 2.491 | 487 | 3896 |
| *Caudate_R* | 18 | 14 | 2 | <0.001 | 5.636 | 3.608 | 469 | 3752 |
| *Caudate_L* | -16 | -18 | 20 | <0.001 | 6.102 | 3.470 | 465 | 3720 |
| *Cuneus_L* | 2 | -74 | 18 | <0.001 | 3.865 | 2.509 | 460 | 3680 |
| *Cuneus_R* | 4 | -74 | 18 | <0.001 | 4.064 | 2.655 | 459 | 3672 |
| *Occipital_Sup_L* | -16 | -78 | 40 | 0.001 | 3.395 | 2.422 | 364 | 2912 |
| *Frontal_Inf_Orb_R* | 44 | 38 | -4 | <0.001 | 3.733 | 2.530 | 361 | 2888 |
| *Rolandic_Oper_L* | -50 | -22 | 14 | <0.001 | 5.017 | 2.800 | 345 | 2760 |
| *Precentral_R* | 64 | 2 | 22 | <0.001 | 4.004 | 2.602 | 329 | 2632 |
| *Frontal_Inf_Orb_L* | -32 | 20 | -18 | <0.001 | 3.817 | 2.529 | 309 | 2472 |
| *Precentral_L* | -34 | -18 | 50 | <0.001 | 4.408 | 2.593 | 305 | 2440 |
| *Occipital_Sup_R* | 22 | -78 | 36 | <0.001 | 4.105 | 2.674 | 275 | 2200 |
| *Precuneus_L* | 0 | -62 | 16 | <0.001 | 4.309 | 2.657 | 268 | 2144 |
| *Parietal_Sup_L* | -28 | -54 | 50 | 0.001 | 3.492 | 2.382 | 242 | 1936 |
| *Frontal_Mid_R* | 32 | 50 | 2 | <0.001 | 3.544 | 2.444 | 235 | 1880 |
| *Hippocampus_L* | -20 | -30 | -4 | <0.001 | 6.352 | 3.160 | 222 | 1776 |
| *Rolandic_Oper_R* | 62 | -4 | 14 | <0.001 | 3.680 | 2.601 | 210 | 1680 |
| *Insula_R* | 28 | 10 | -16 | <0.001 | 3.689 | 2.592 | 201 | 1608 |
| *SupraMarginal_R* | 40 | -34 | 38 | <0.001 | 3.987 | 2.685 | 188 | 1504 |
| *Heschl_R* | 38 | -24 | 14 | <0.001 | 3.817 | 2.725 | 169 | 1352 |
| *Amygdala_R* | 32 | 0 | -18 | <0.001 | 6.812 | 4.009 | 154 | 1232 |
| *Insula_L* | -26 | 8 | -14 | <0.001 | 5.338 | 3.102 | 149 | 1192 |
| *Frontal_Sup_L* | -18 | 2 | 56 | 0.003 | 3.055 | 2.411 | 147 | 1176 |
| *Pallidum_R* | 18 | 8 | 0 | <0.001 | 5.022 | 3.197 | 145 | 1160 |
| *Heschl_L* | -54 | -10 | 8 | <0.001 | 5.106 | 3.282 | 142 | 1136 |
| *Occipital_Inf_R* | 20 | -92 | -6 | <0.001 | 4.471 | 2.611 | 139 | 1112 |
| *Fusiform_L* | -22 | -46 | -12 | 0.001 | 3.440 | 2.592 | 136 | 1088 |
| *Temporal_Pole_Sup_L* | -34 | 4 | -20 | <0.001 | 5.790 | 3.130 | 124 | 992 |
| *Frontal_Inf_Tri_L* | -48 | 16 | 6 | 0.007 | 2.789 | 2.236 | 122 | 976 |
| *Parietal_Inf_R* | 40 | -40 | 40 | 0.005 | 2.943 | 2.313 | 103 | 824 |
| *Hippocampus_R* | 18 | -30 | -4 | <0.001 | 7.510 | 3.762 | 99 | 792 |
| *SupraMarginal_L* | -52 | -24 | 14 | <0.001 | 3.613 | 2.426 | 98 | 784 |
| *ParaHippocampal_R* | 22 | 6 | -20 | <0.001 | 4.637 | 2.739 | 91 | 728 |
| *Amygdala_L* | -30 | 2 | -18 | <0.001 | 5.688 | 3.446 | 86 | 688 |
| *Olfactory_R* | 22 | 8 | -14 | <0.001 | 5.564 | 3.582 | 69 | 552 |
| *ParaHippocampal_L* | -18 | -36 | -4 | <0.001 | 3.650 | 2.635 | 58 | 464 |
| *Occipital_Mid_R* | 32 | -70 | 28 | 0.010 | 2.530 | 2.190 | 57 | 456 |
| *Frontal_Sup_R* | 32 | 64 | 10 | 0.010 | 2.622 | 2.246 | 54 | 432 |
| *Frontal_Sup_Orb_L* | -24 | 10 | -14 | <0.001 | 5.237 | 2.960 | 51 | 408 |
| *Frontal_Mid_Orb_R* | 48 | 50 | -2 | 0.004 | 3.070 | 2.371 | 50 | 400 |

**Table S4.** **Anatomical characterization of atrophied regions in the 2nd Cohort.** We only represented those regions having more than 50 voxels of overlapping between with the map of volume loss, resulting from the group comparison contrast DM1 < HC.

| *Area (AAL)* | *Peak x (MNI)* | *Peak y (MNI)* | *Peak z (MNI)* | *Peak p* | *Peak t* | *Mean t* | *N Voxels* | *Vol (mm<sup>3</sup>)* |
| --- | --- | --- | --- | --- | --- | --- | --- | --- |
| *Cerebellum* | -10 | -82 | -34 | <0.001 | 4.895 | 2.787 | 6780 | 54240 |
| *Postcentral_L* | -56 | -4 | 44 | <0.001 | 4.798 | 2.867 | 1455 | 11640 |
| *Thalamus_L* | -18 | -24 | 12 | <0.001 | 8.675 | 5.260 | 1043 | 8344 |
| *Thalamus_R* | 18 | -24 | 12 | <0.001 | 8.665 | 5.241 | 987 | 7896 |
| *Frontal_Inf_Tri_R* | 58 | 38 | 16 | <0.001 | 4.390 | 2.610 | 956 | 7648 |
| *Parietal_Inf_L* | -42 | -42 | 42 | <0.001 | 6.180 | 3.218 | 950 | 7600 |
| *Lingual_L* | -28 | -96 | -16 | <0.001 | 4.881 | 2.901 | 936 | 7488 |
| *Temporal_Sup_R* | 60 | -6 | -6 | <0.001 | 3.718 | 2.522 | 925 | 7400 |
| *Frontal_Inf_Tri_L* | -42 | 26 | 0 | <0.001 | 4.444 | 2.877 | 907 | 7256 |
| *Putamen_L* | -24 | 16 | 2 | <0.001 | 8.163 | 4.912 | 876 | 7008 |
| *Postcentral_R* | 50 | -12 | 28 | <0.001 | 5.501 | 2.924 | 859 | 6872 |
| *Caudate_L* | -14 | 16 | -2 | <0.001 | 6.778 | 4.377 | 825 | 6600 |
| *Putamen_R* | 20 | 14 | 2 | <0.001 | 7.007 | 3.948 | 775 | 6200 |
| *Lingual_R* | 20 | -94 | -12 | <0.001 | 4.435 | 2.932 | 774 | 6192 |
| *Occipital_Mid_L* | -32 | -90 | -6 | <0.001 | 5.038 | 2.761 | 730 | 5840 |
| *Caudate_R* | 18 | 14 | 2 | <0.001 | 6.908 | 4.149 | 710 | 5680 |
| *Parietal_Sup_R* | 30 | -50 | 60 | <0.001 | 4.699 | 2.691 | 604 | 4832 |
| *Precentral_R* | 46 | -12 | 38 | <0.001 | 5.030 | 2.940 | 598 | 4784 |
| *Calcarine_R* | 24 | -98 | 2 | <0.001 | 4.061 | 2.851 | 573 | 4584 |
| *Temporal_Sup_L* | -52 | -30 | 22 | <0.001 | 4.036 | 2.769 | 547 | 4376 |
| *Precentral_L* | -58 | -2 | 42 | <0.001 | 4.614 | 2.827 | 542 | 4336 |
| *Calcarine_L* | -8 | -60 | 10 | <0.001 | 4.988 | 2.800 | 536 | 4288 |
| *Fusiform_L* | -30 | -74 | -14 | <0.001 | 4.472 | 2.998 | 511 | 4088 |
| *Occipital_Inf_L* | -32 | -88 | -6 | <0.001 | 5.138 | 3.100 | 482 | 3856 |
| *Parietal_Inf_R* | 38 | -50 | 54 | <0.001 | 3.936 | 2.709 | 482 | 3856 |
| *Occipital_Inf_R* | 32 | -92 | -2 | <0.001 | 4.685 | 3.008 | 469 | 3752 |
| *SupraMarginal_L* | -50 | -26 | 24 | <0.001 | 4.972 | 2.789 | 416 | 3328 |
| *Frontal_Inf_Oper_R* | 50 | 8 | 16 | 0.001 | 3.494 | 2.458 | 380 | 3040 |
| *Precuneus_R* | 14 | -50 | 20 | <0.001 | 3.743 | 2.478 | 369 | 2952 |
| *Occipital_Mid_R* | 32 | -96 | 6 | <0.001 | 4.929 | 3.360 | 353 | 2824 |
| *Frontal_Inf_Orb_L* | -40 | 32 | -6 | <0.001 | 4.467 | 3.000 | 325 | 2600 |
| *Insula_L* | -28 | 16 | 4 | <0.001 | 4.972 | 2.615 | 325 | 2600 |
| *Cingulum_Mid_R* | 10 | -34 | 34 | 0.001 | 3.478 | 2.573 | 296 | 2368 |
| *Cingulum_Mid_L* | -2 | -34 | 36 | <0.001 | 3.549 | 2.589 | 254 | 2032 |
| *Rolandic_Oper_L* | -48 | -26 | 22 | <0.001 | 4.179 | 2.904 | 218 | 1744 |
| *Precuneus_L* | -8 | -58 | 10 | <0.001 | 4.892 | 2.978 | 189 | 1512 |
| *SupraMarginal_R* | 48 | -16 | 28 | <0.001 | 3.573 | 2.329 | 183 | 1464 |
| *Frontal_Inf_Orb_R* | 36 | 30 | -4 | <0.001 | 3.556 | 2.601 | 182 | 1456 |
| *Parietal_Sup_L* | -40 | -42 | 56 | <0.001 | 4.560 | 2.778 | 169 | 1352 |
| *Occipital_Sup_L* | -16 | -88 | 28 | 0.001 | 3.441 | 2.530 | 162 | 1296 |
| *Pallidum_R* | 20 | 8 | 2 | <0.001 | 5.754 | 3.467 | 149 | 1192 |
| *Pallidum_L* | -18 | 6 | 4 | <0.001 | 5.938 | 3.369 | 122 | 976 |
| *Occipital_Sup_R* | 26 | -96 | 10 | <0.001 | 4.176 | 3.110 | 120 | 960 |
| *Rolandic_Oper_R* | 58 | 2 | 4 | 0.003 | 3.115 | 2.452 | 119 | 952 |
| *Insula_R* | 32 | 28 | -2 | <0.001 | 3.992 | 2.747 | 110 | 880 |
| *Angular_R* | 38 | -58 | 52 | <0.001 | 3.615 | 2.575 | 110 | 880 |
| *Cingulum_Post_R* | 14 | -48 | 20 | 0.005 | 2.955 | 2.294 | 104 | 832 |
| *Temporal_Mid_R* | 50 | -14 | -14 | <0.001 | 3.962 | 2.802 | 104 | 832 |
| *Amygdala_R* | 24 | 2 | -12 | <0.001 | 4.594 | 2.648 | 103 | 824 |
| *Cuneus_R* | 22 | -96 | 10 | <0.001 | 3.552 | 2.647 | 99 | 792 |
| *Hippocampus_L* | -20 | -28 | -6 | <0.001 | 4.259 | 2.914 | 91 | 728 |
| *Fusiform_R* | 30 | -84 | -4 | <0.001 | 4.219 | 2.408 | 90 | 720 |
| *Temporal_Mid_L* | -58 | -32 | 8 | 0.004 | 2.997 | 2.316 | 83 | 664 |
| *Frontal_Mid_R* | 44 | 32 | 18 | <0.001 | 3.627 | 2.527 | 75 | 600 |
| *Olfactory_L* | -24 | 6 | -14 | <0.001 | 3.924 | 2.741 | 60 | 480 |
| *Amygdala_L* | -18 | 0 | -12 | <0.001 | 3.996 | 2.716 | 60 | 480 |
| *Temporal_Pole_Sup_R* | 58 | 2 | 2 | 0.001 | 3.326 | 2.522 | 56 | 448 |
| *Cingulum_Post_L* | -2 | -34 | 32 | 0.003 | 3.126 | 2.499 | 55 | 440 |
| *Heschl_L* | -36 | -30 | 14 | 0.004 | 3.079 | 2.420 | 54 | 432 |
| *Olfactory_R* | 6 | 18 | -4 | <0.001 | 4.663 | 2.860 | 50 | 400 |

**Table S5.**
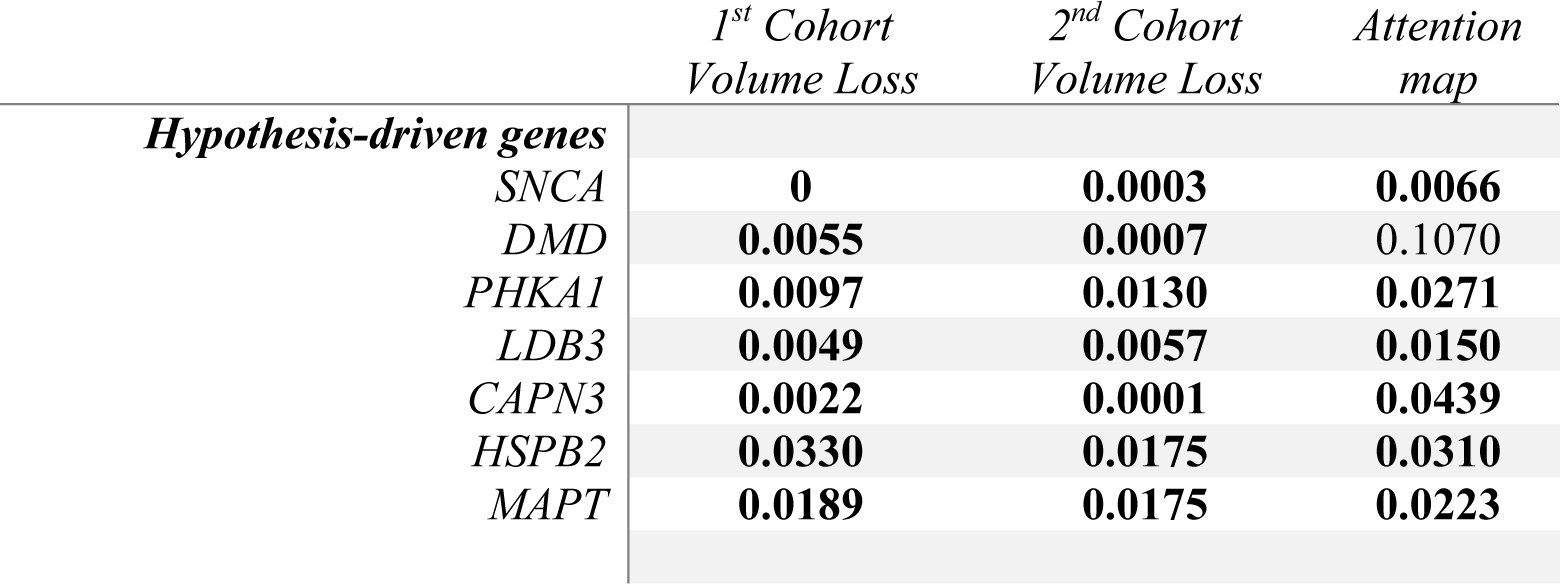
Statistical significance (p-value) of hypothesis-driven genes using surrogate-data.

**Table S6.**
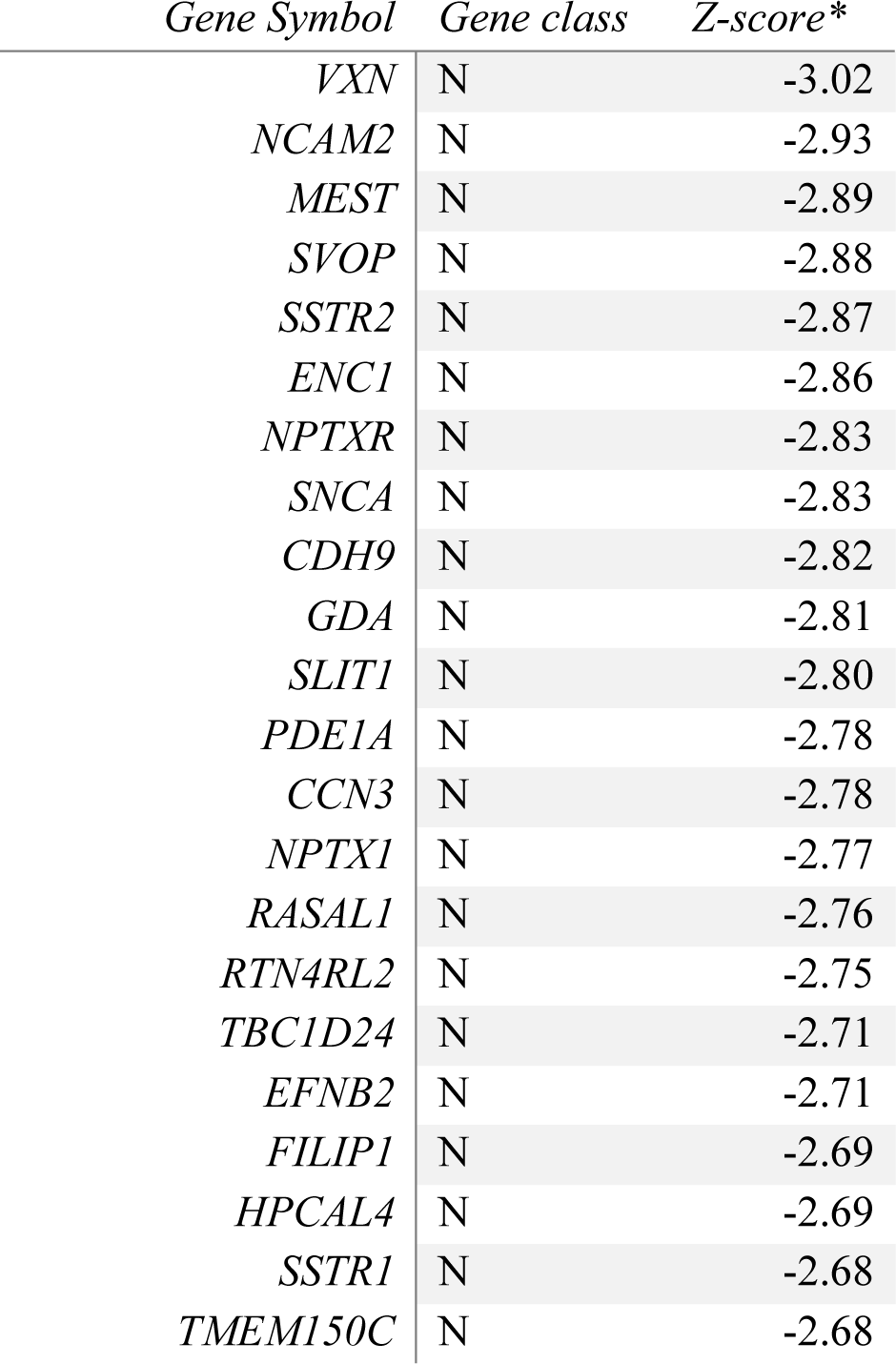
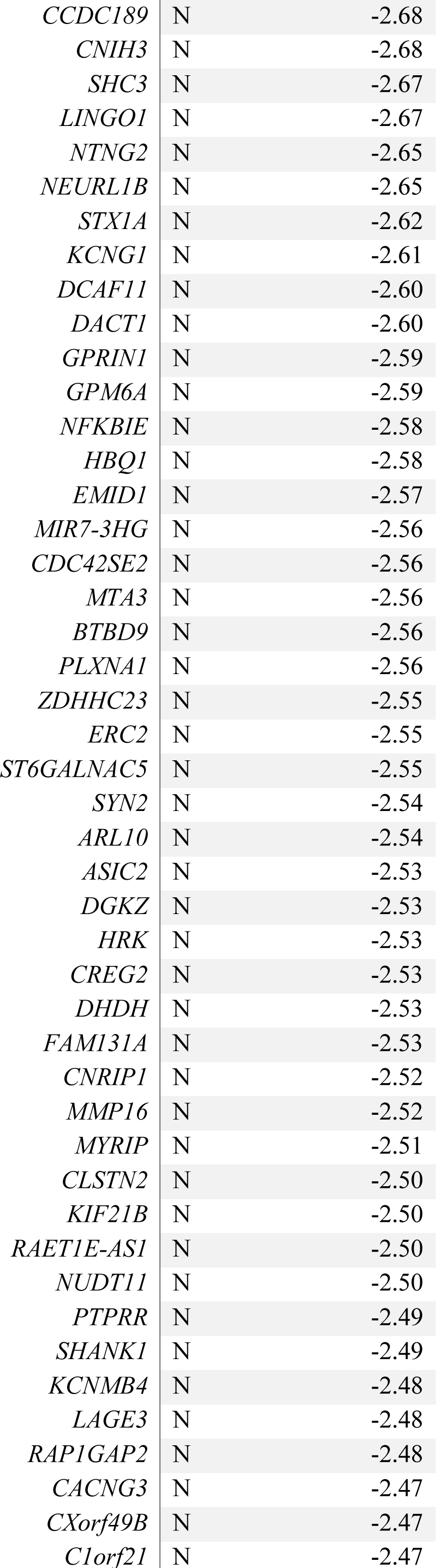
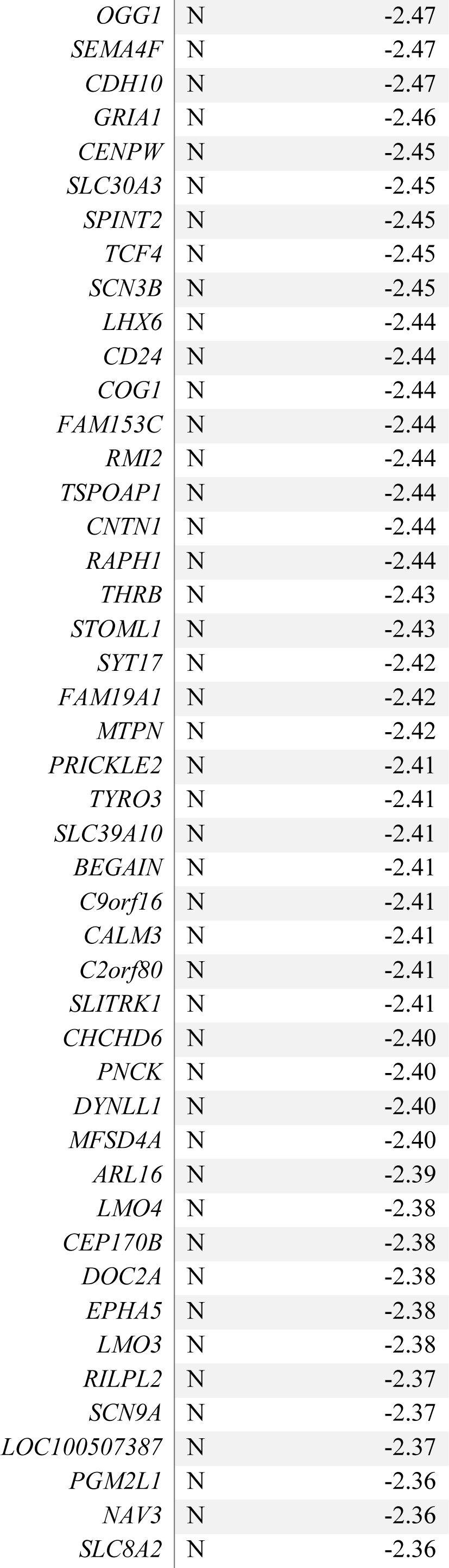
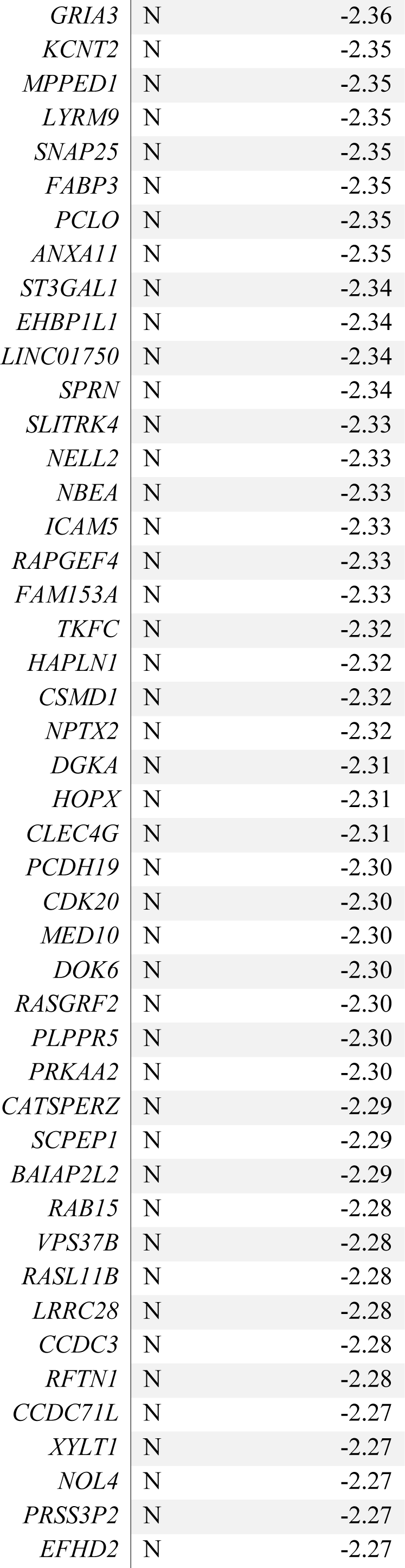
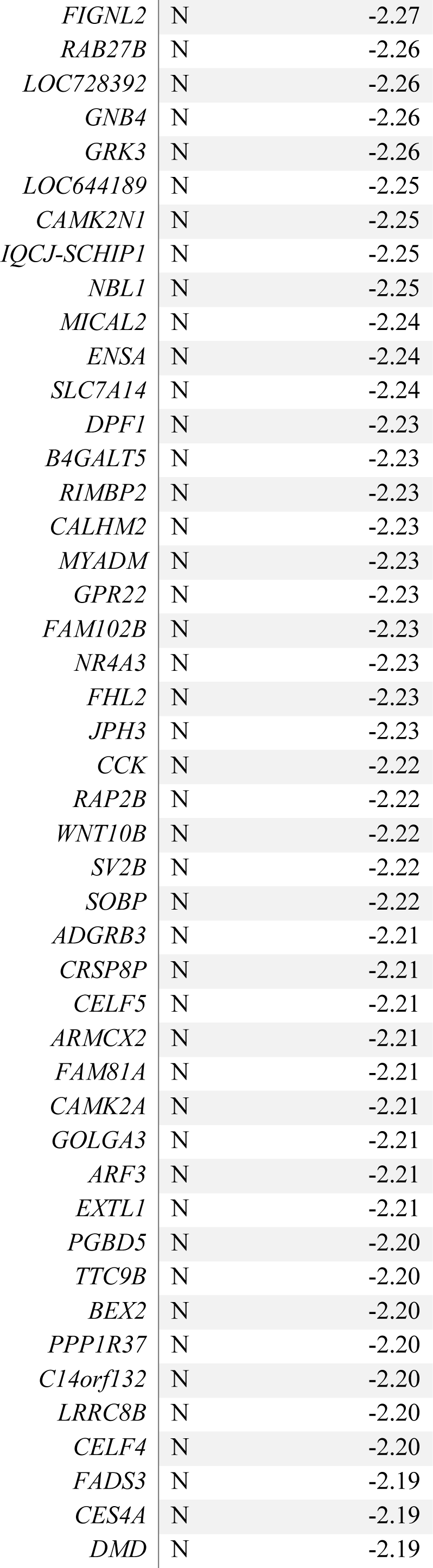
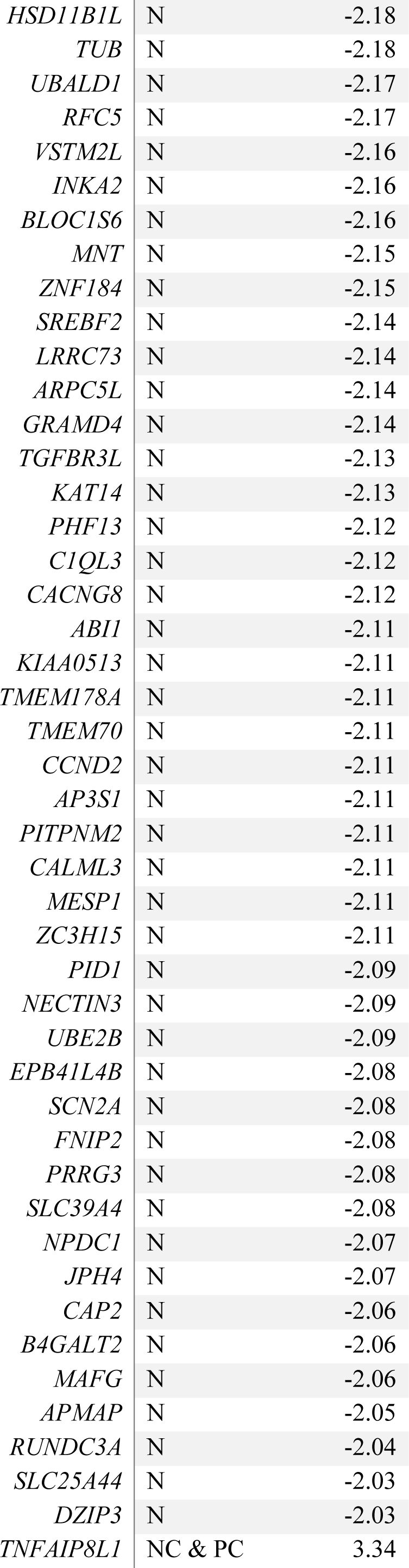
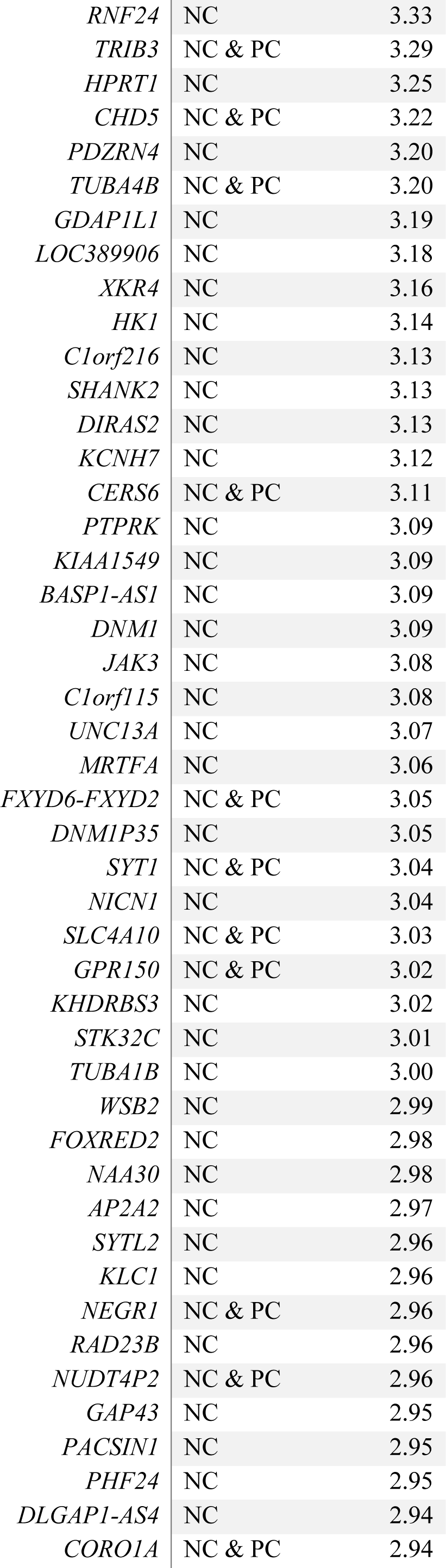
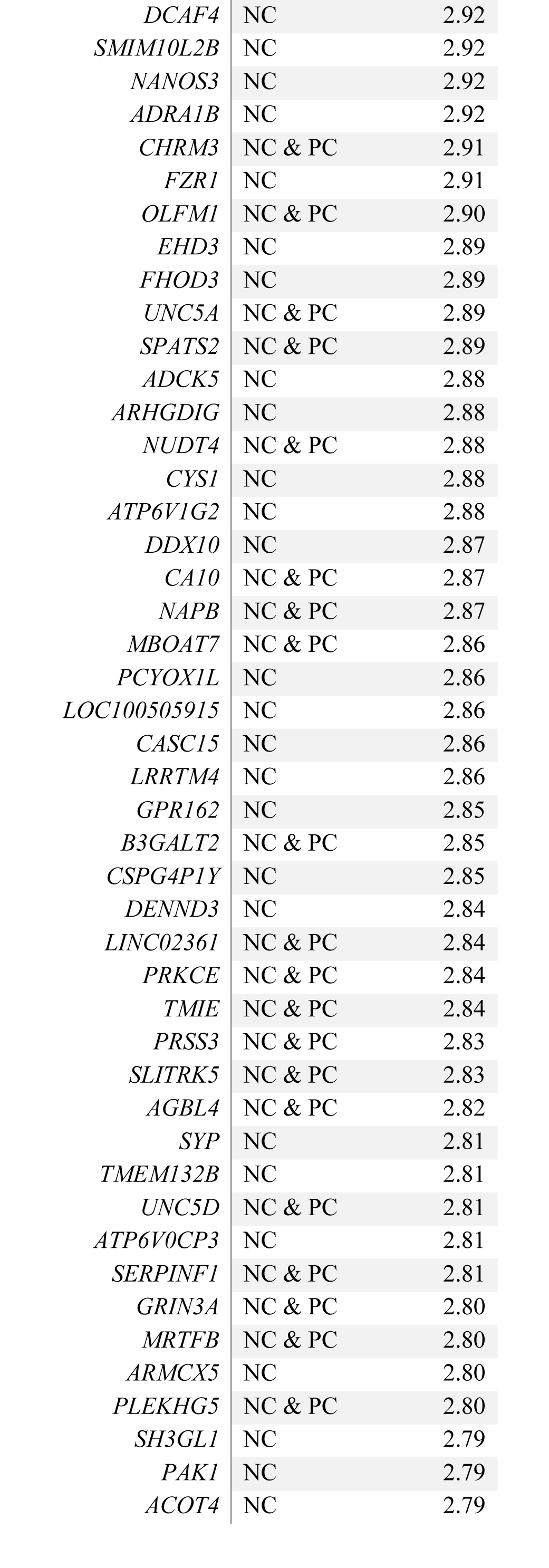
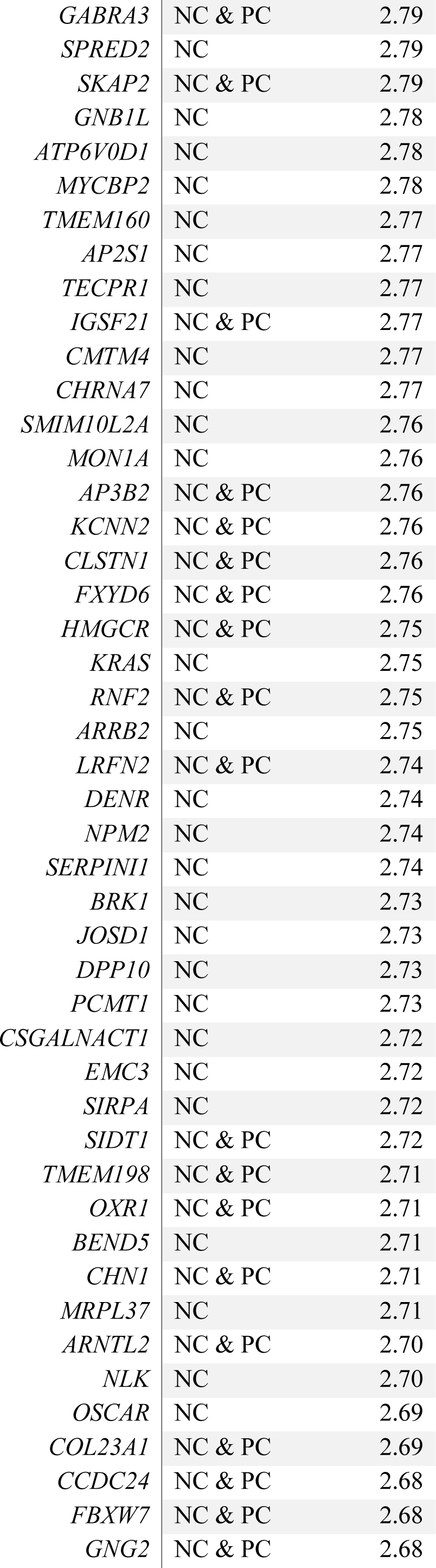
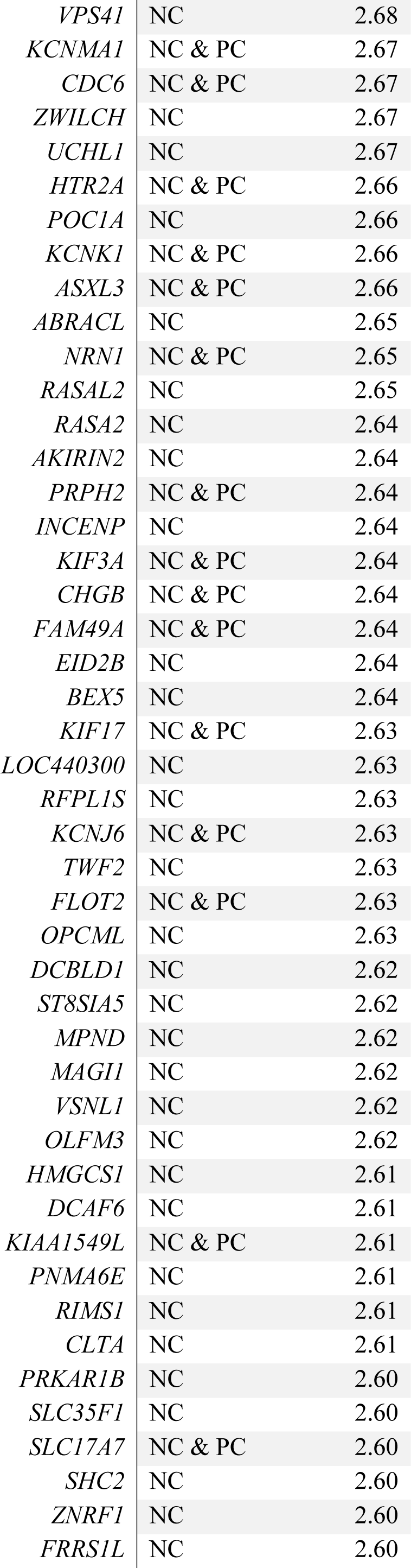
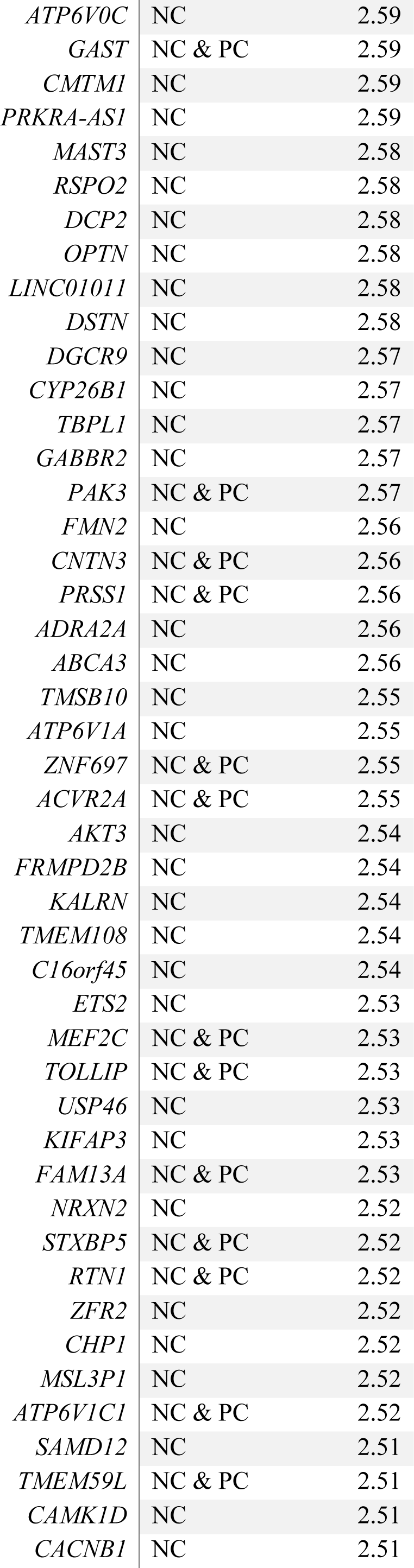
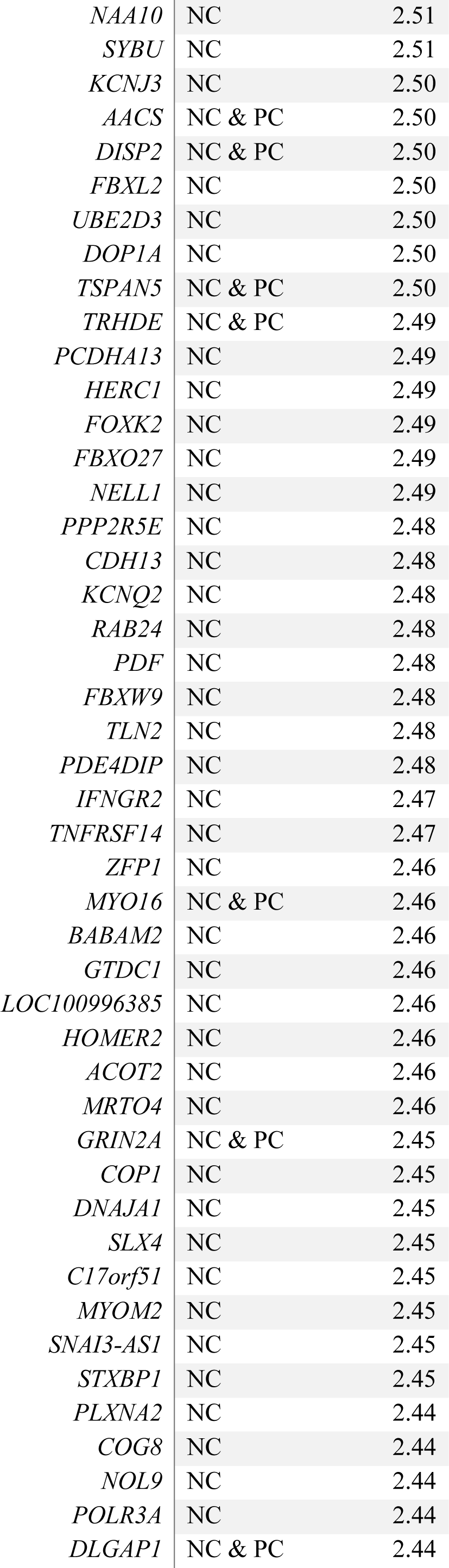
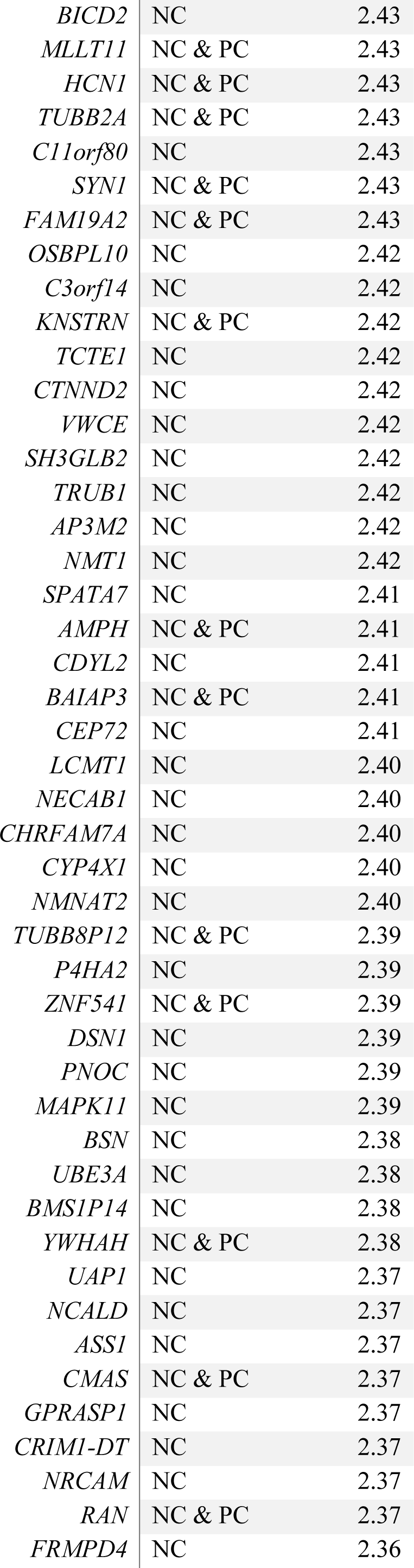
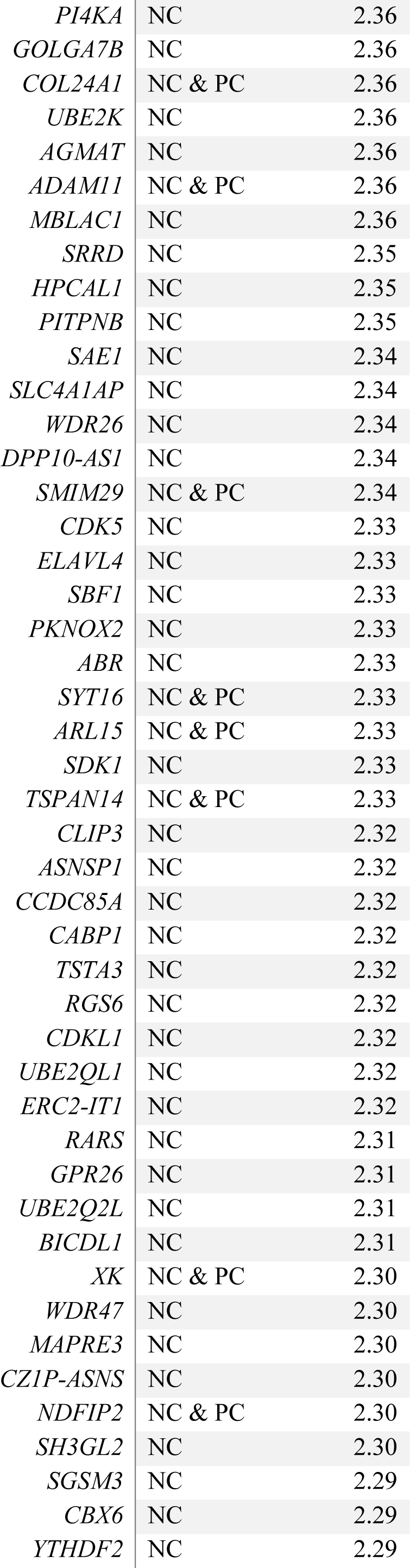
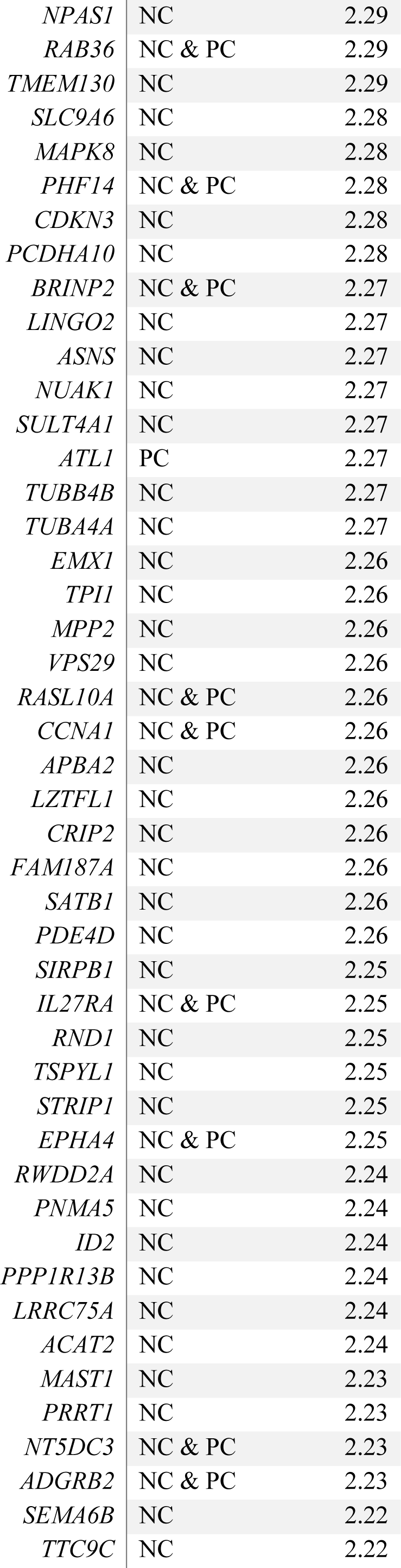
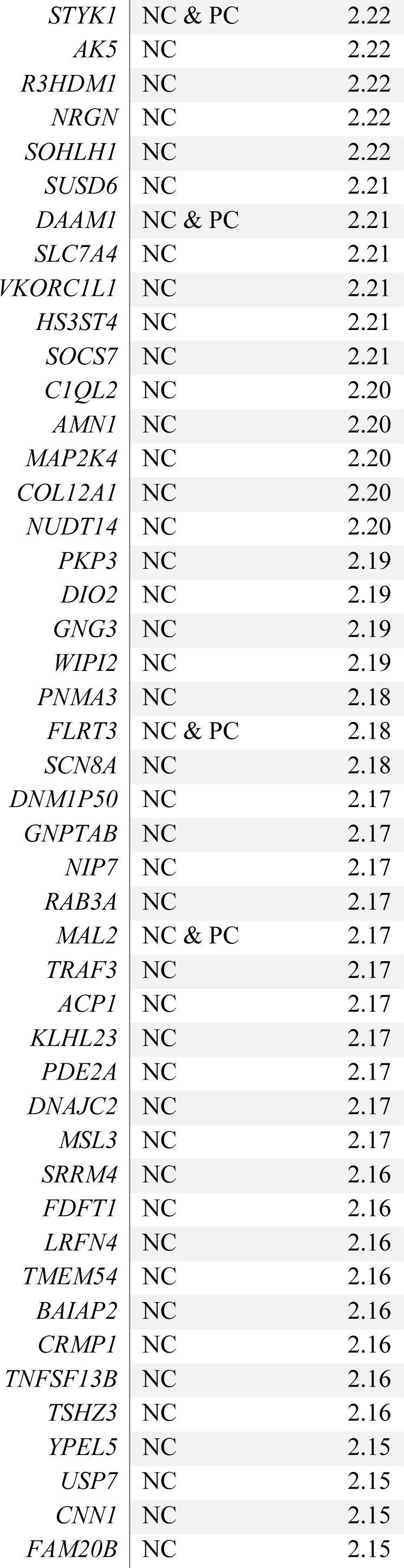
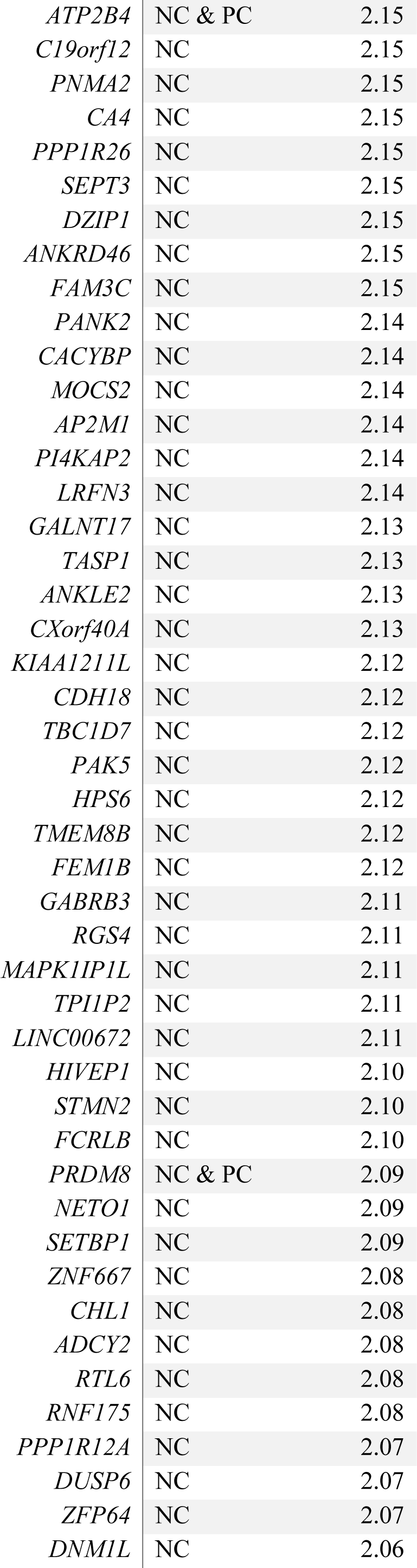
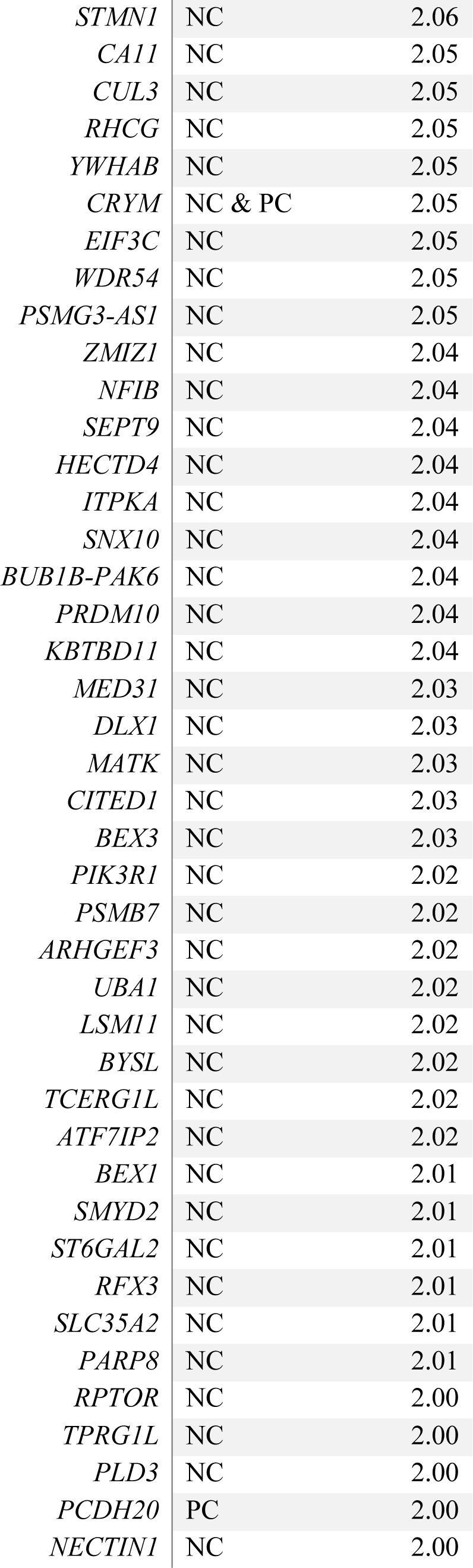
**Negative-correlated genes (blue-cluster) used for enrichment (Volume Loss).** Z-score* represents for N genes the mean Z-score over the two cohorts in the probability distribution of the spatial correlation between gene-transcription activity and volume loss maps; for connector (C) hubs genes it represents the Z-score in the probability distribution of strength values towards the N-genes cluster.

**Table S7.**
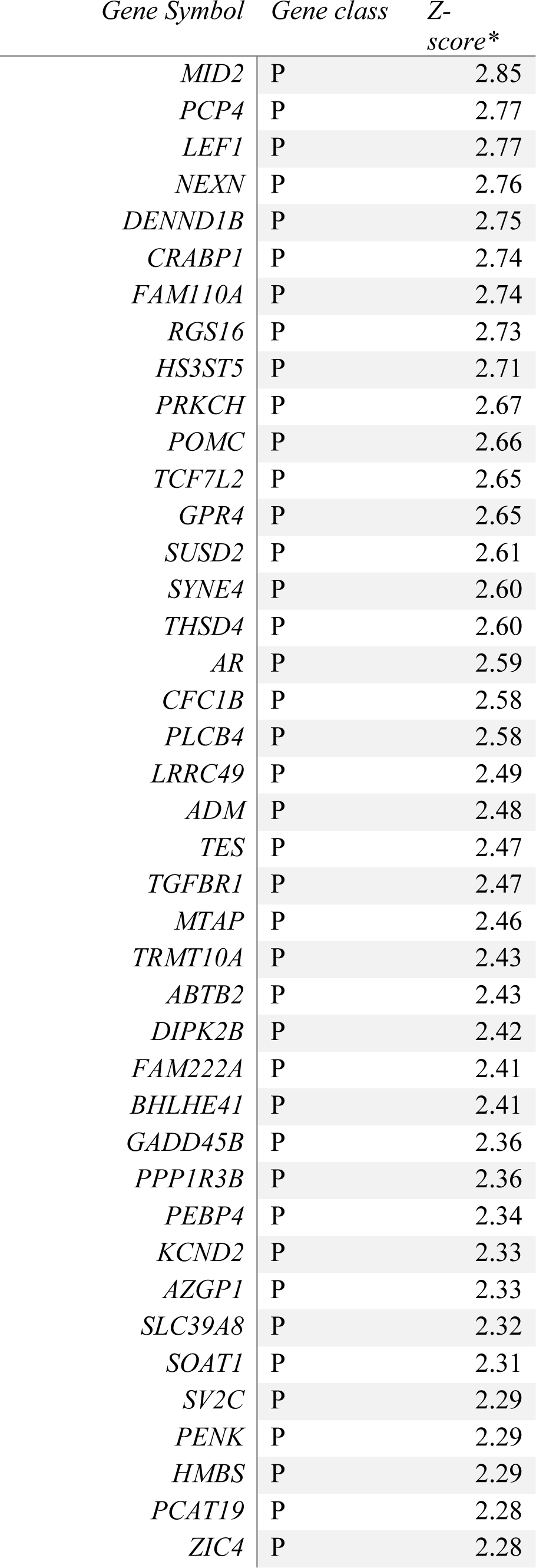
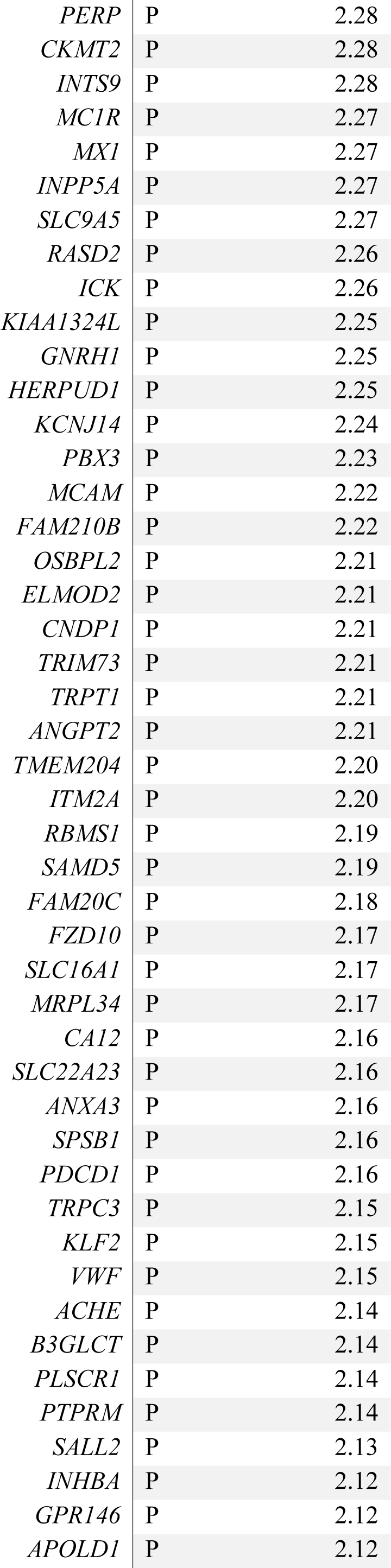
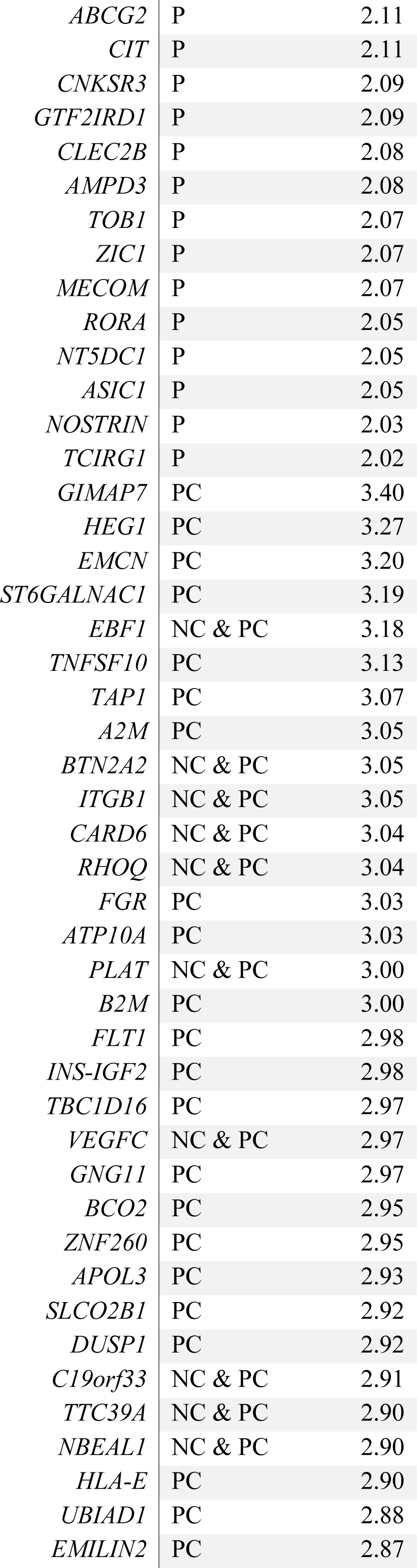
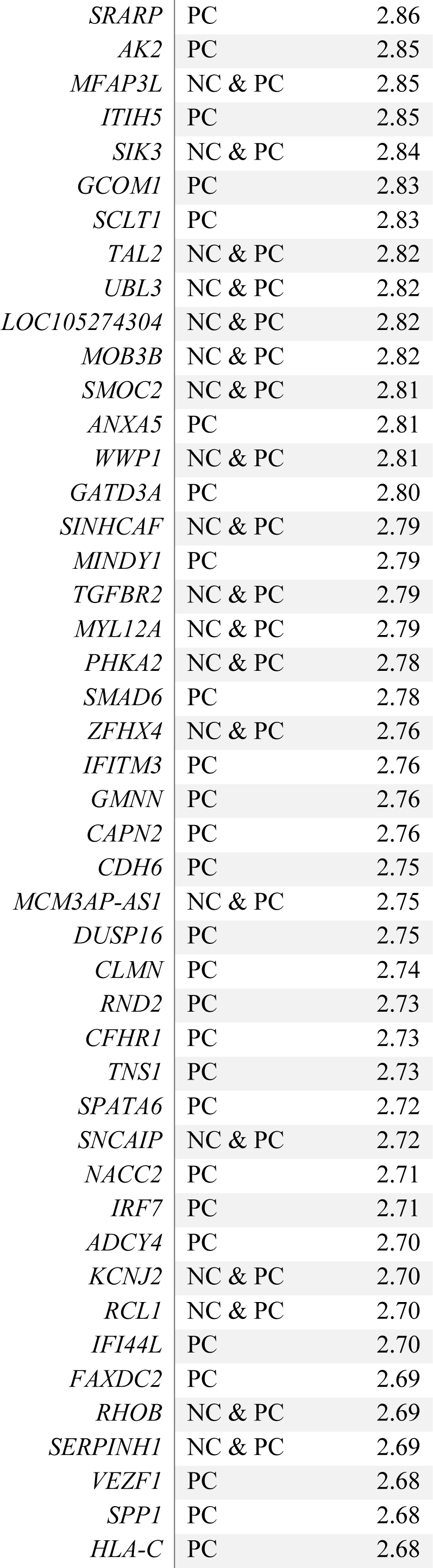
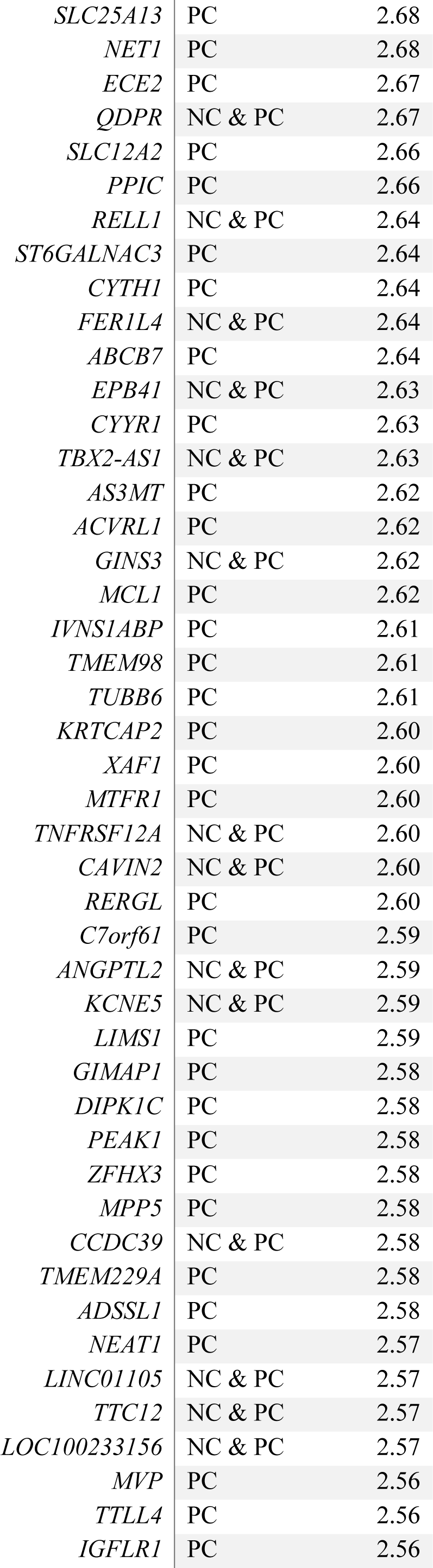
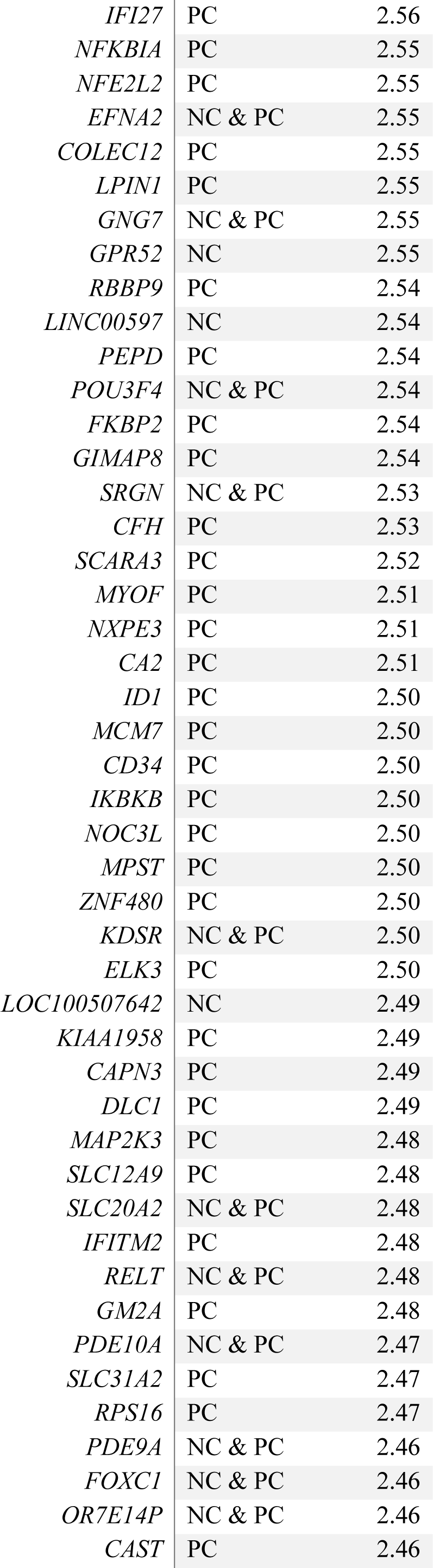
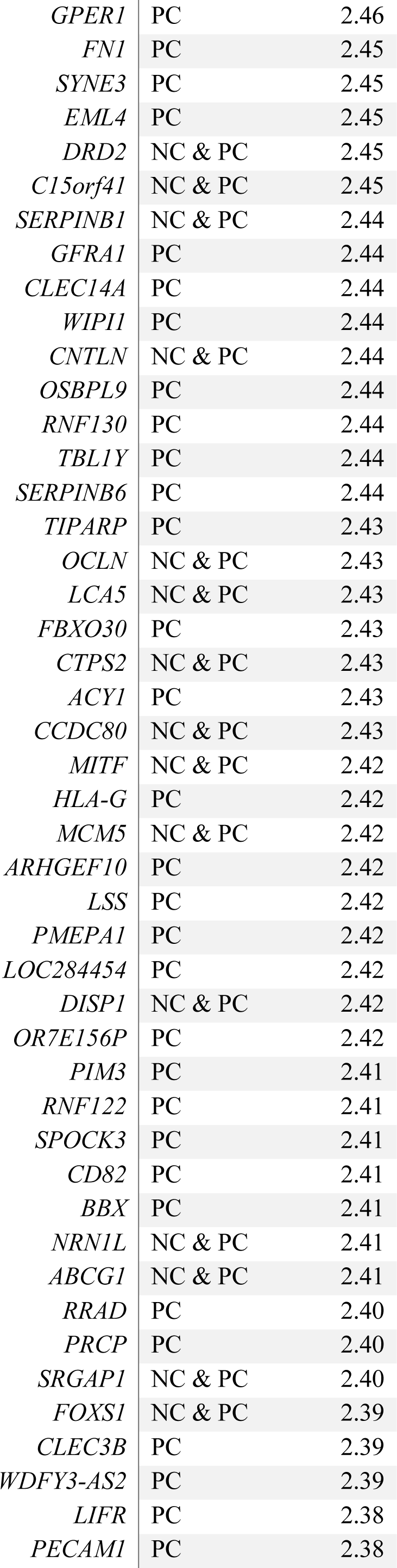
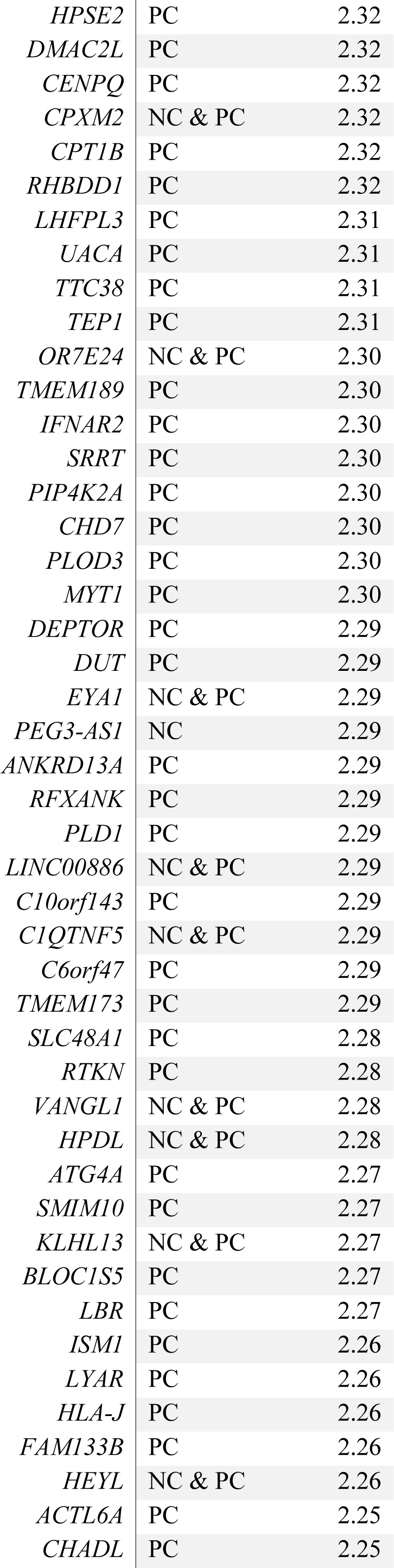
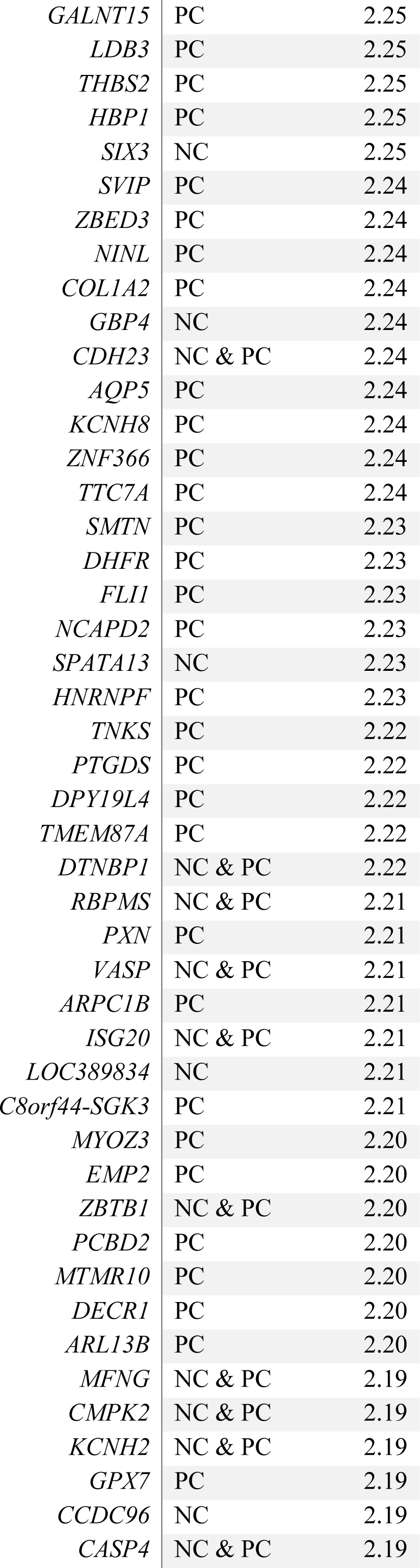
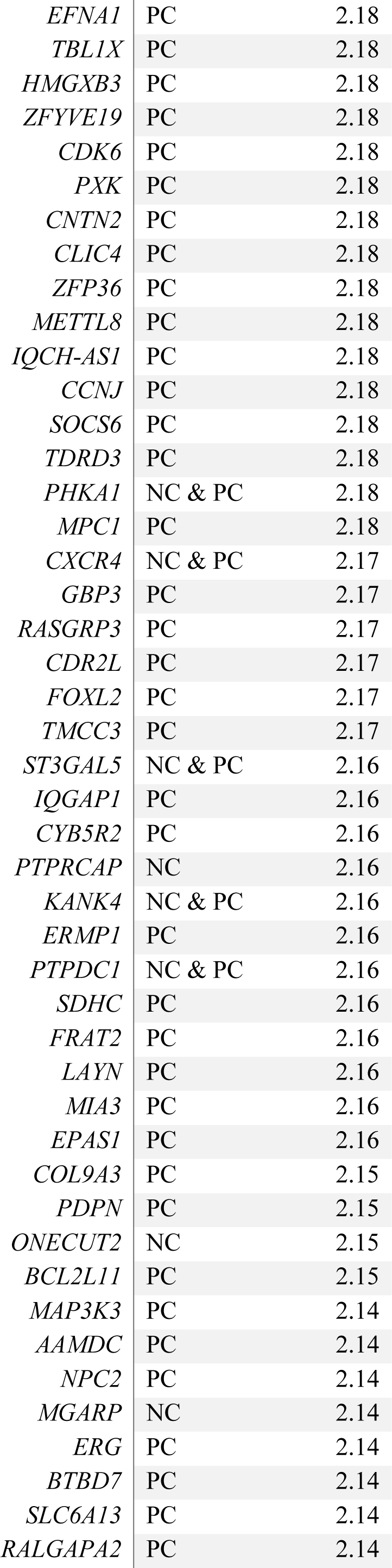
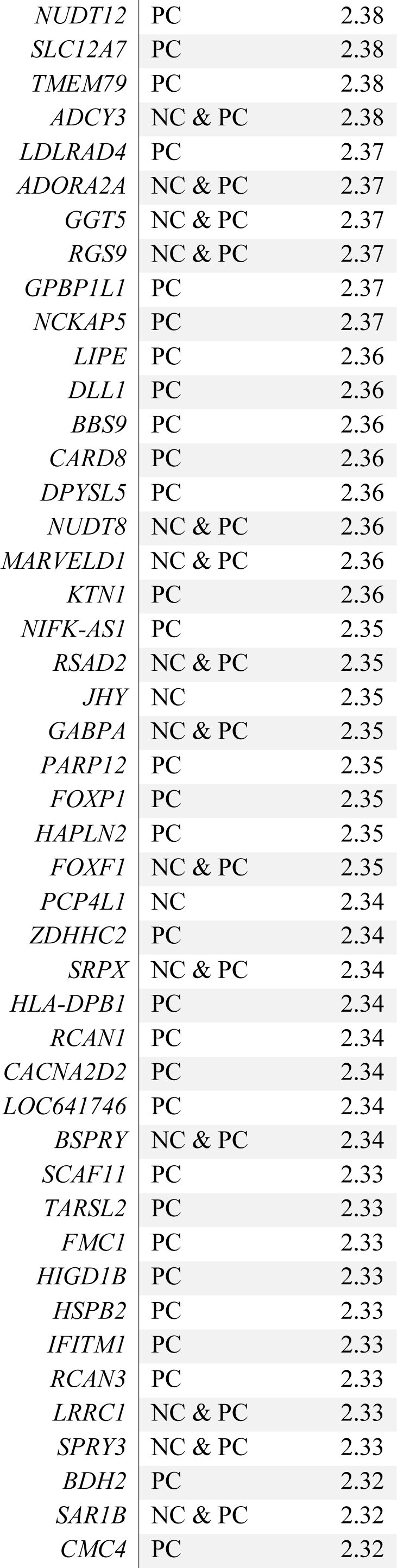
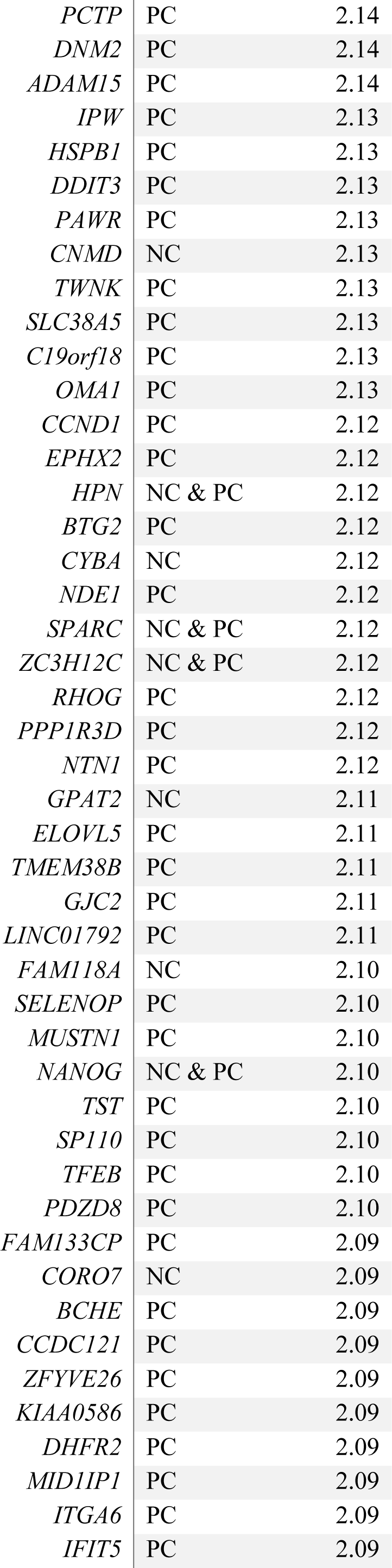
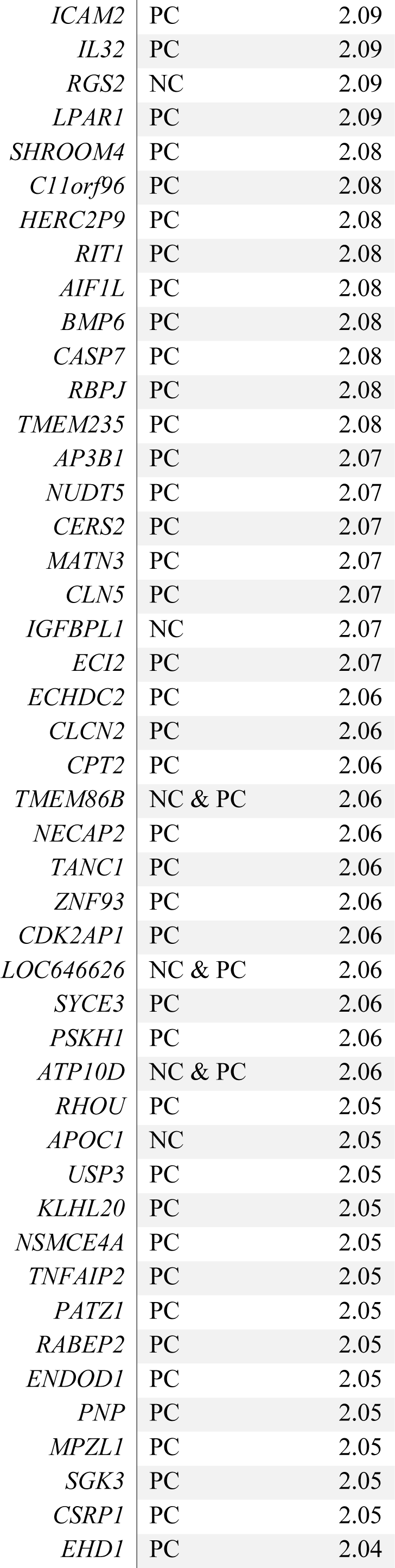
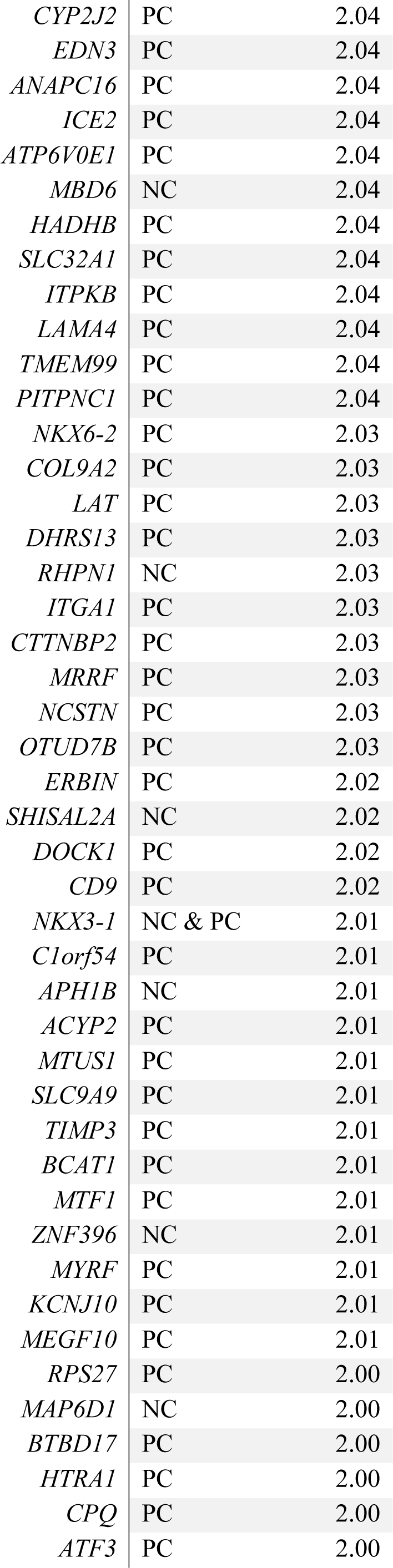
**Positive-correlated genes (red-cluster) used for enrichment (Volume Loss).** Z-score* represents for P genes the mean Z-score over the two cohorts in the probability distribution of the spatial correlation between gene-transcription activity and volume loss maps; for connector (C) hubs genes it represents the Z-score in the probability distribution of strength values towards the P-genes cluster.

**Table S8.**
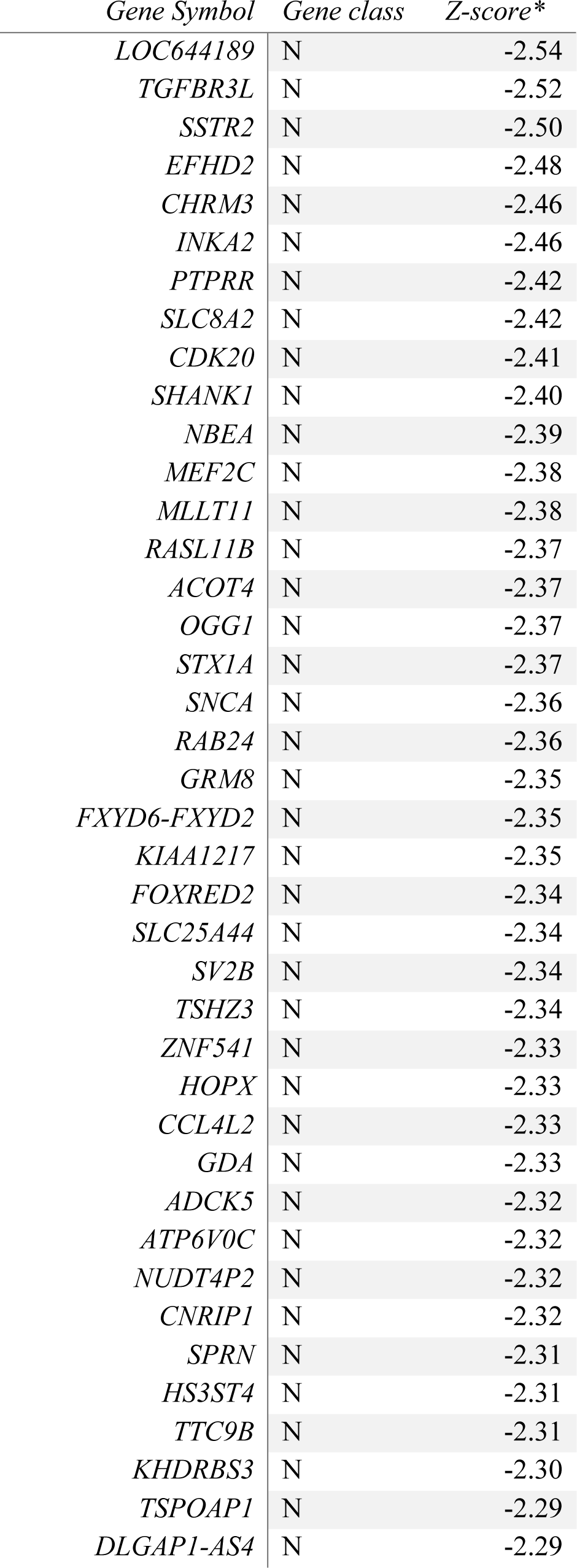
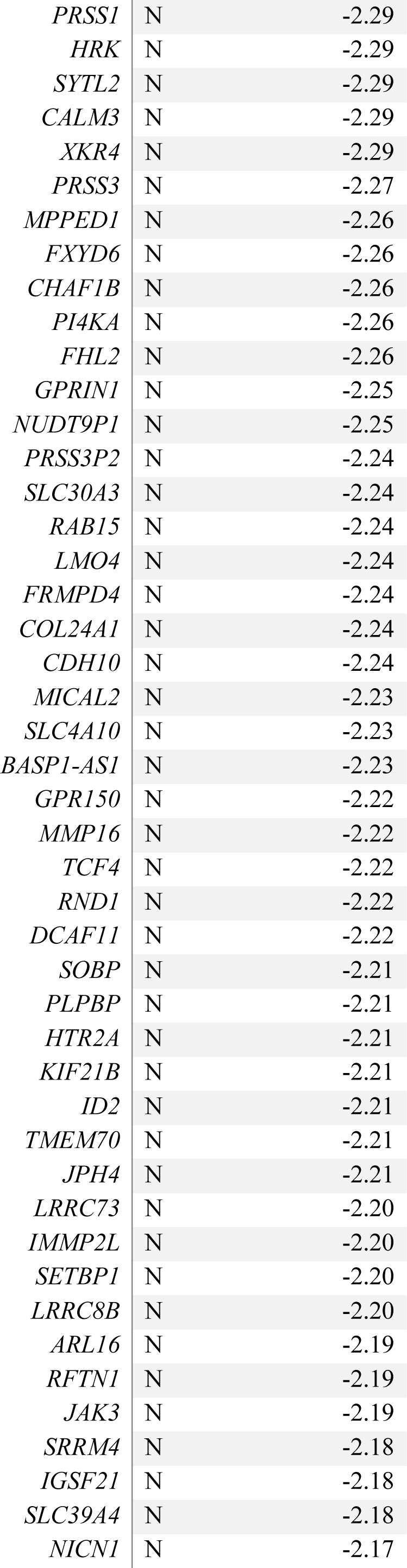
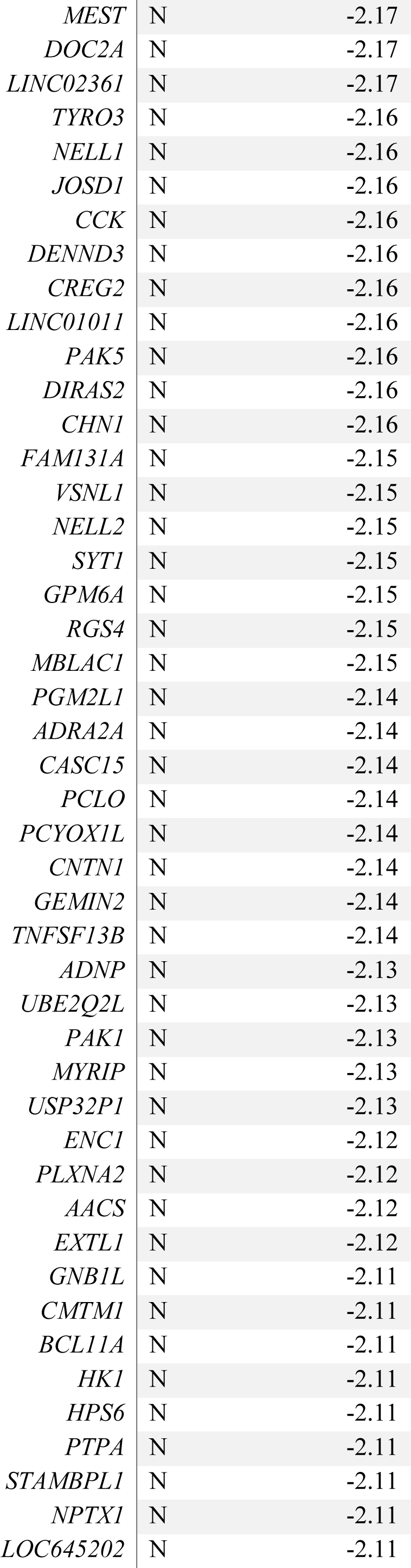
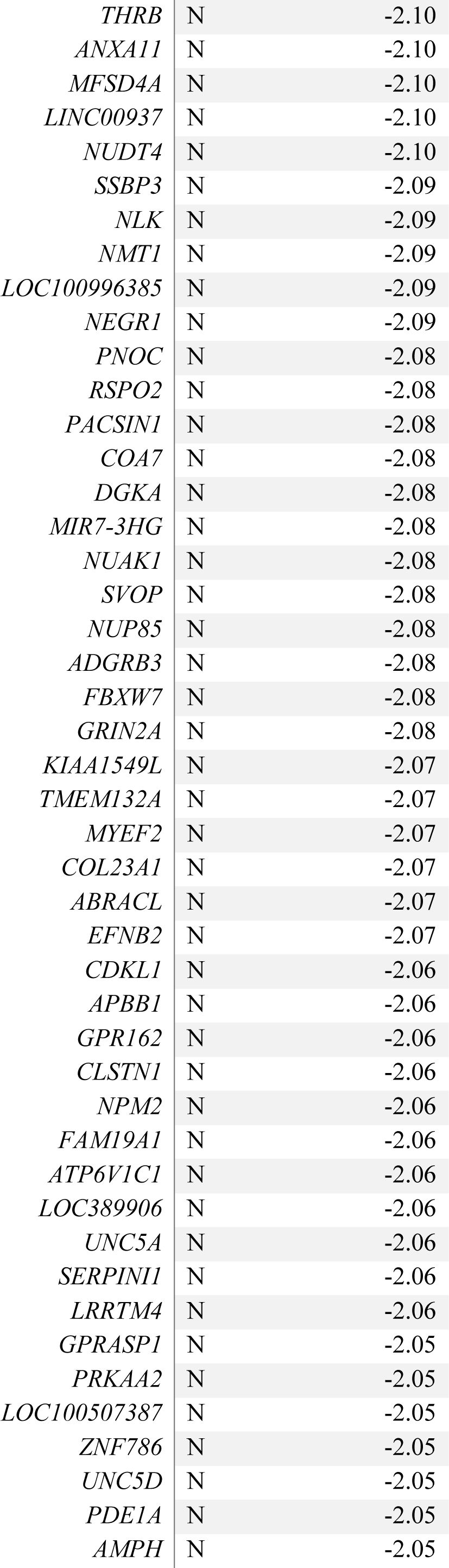
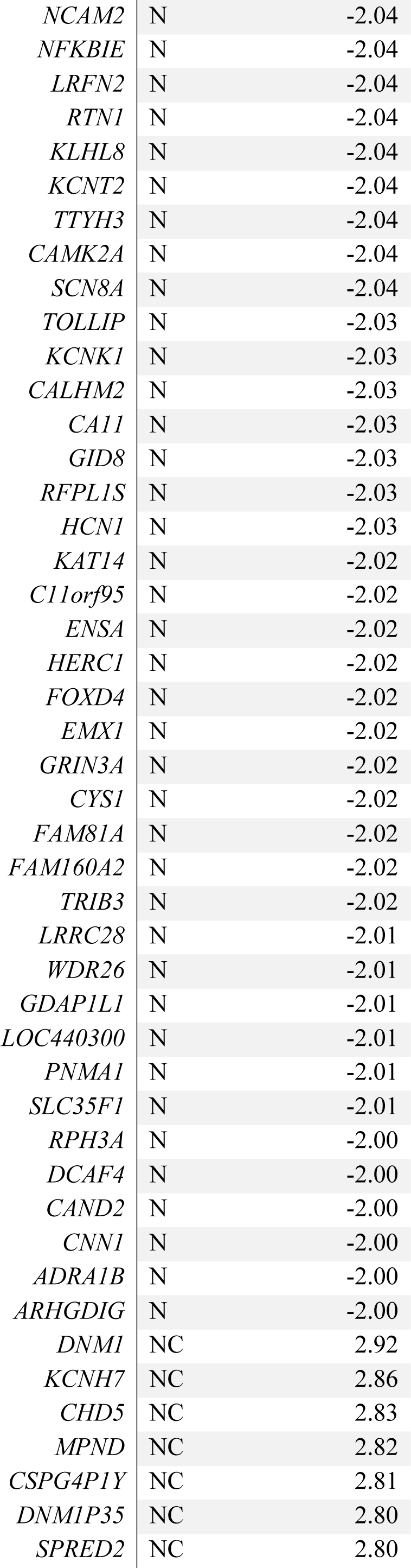
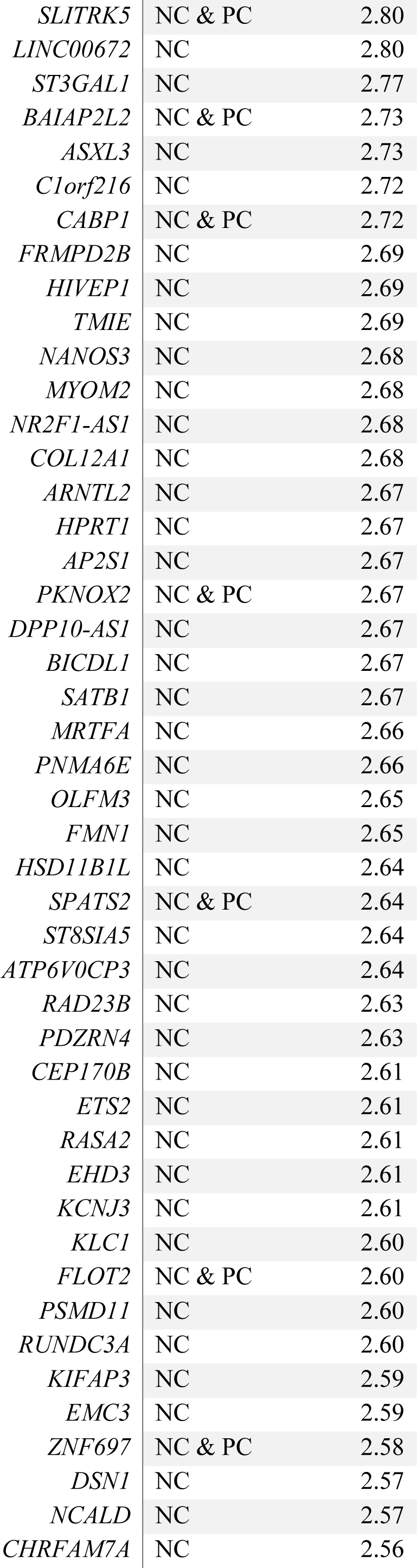
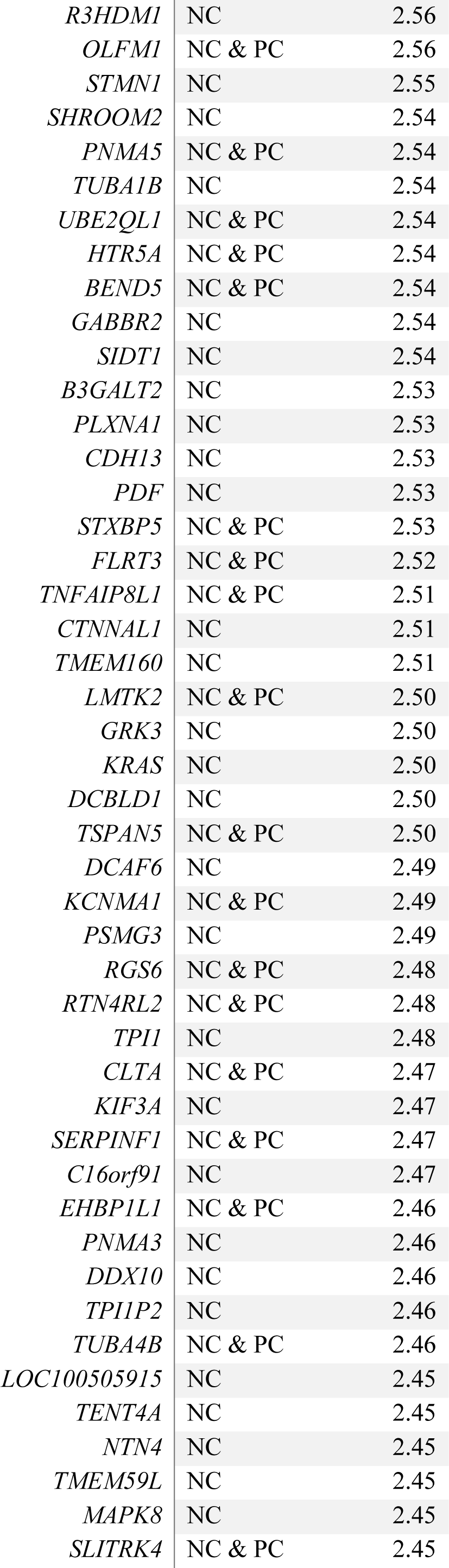
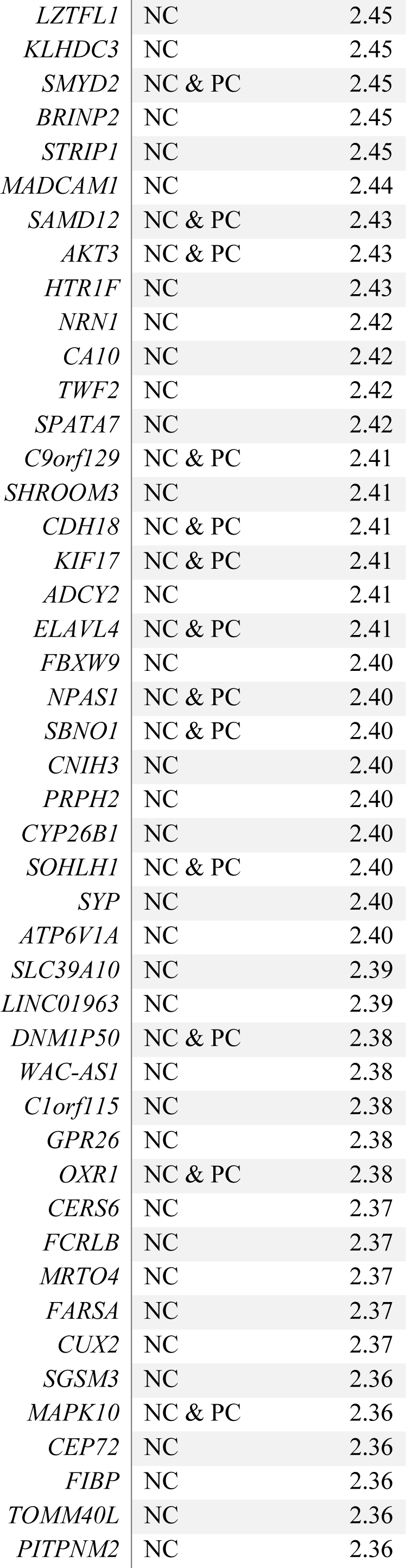
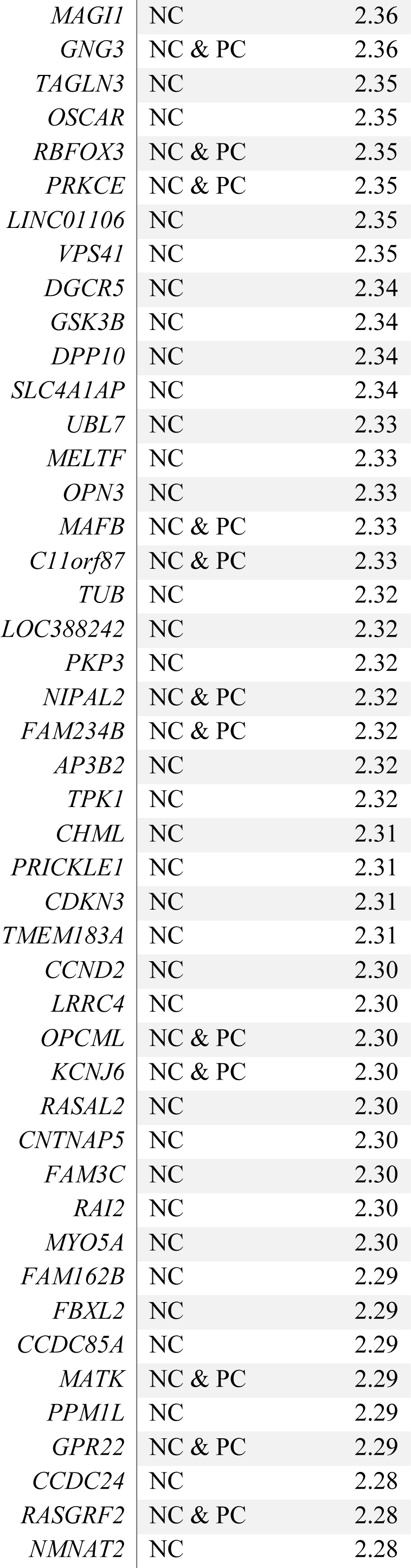
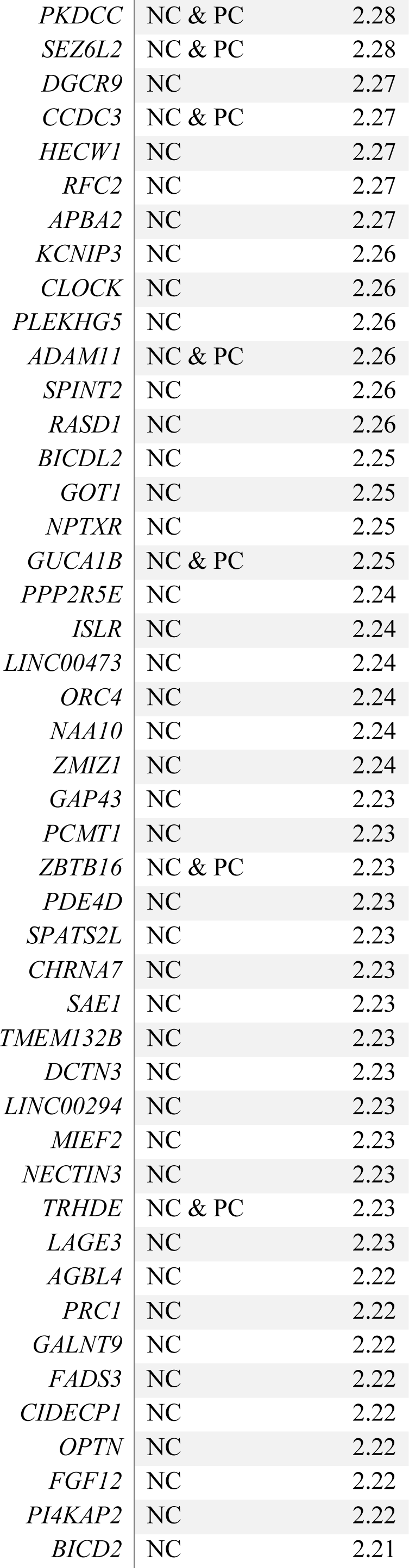
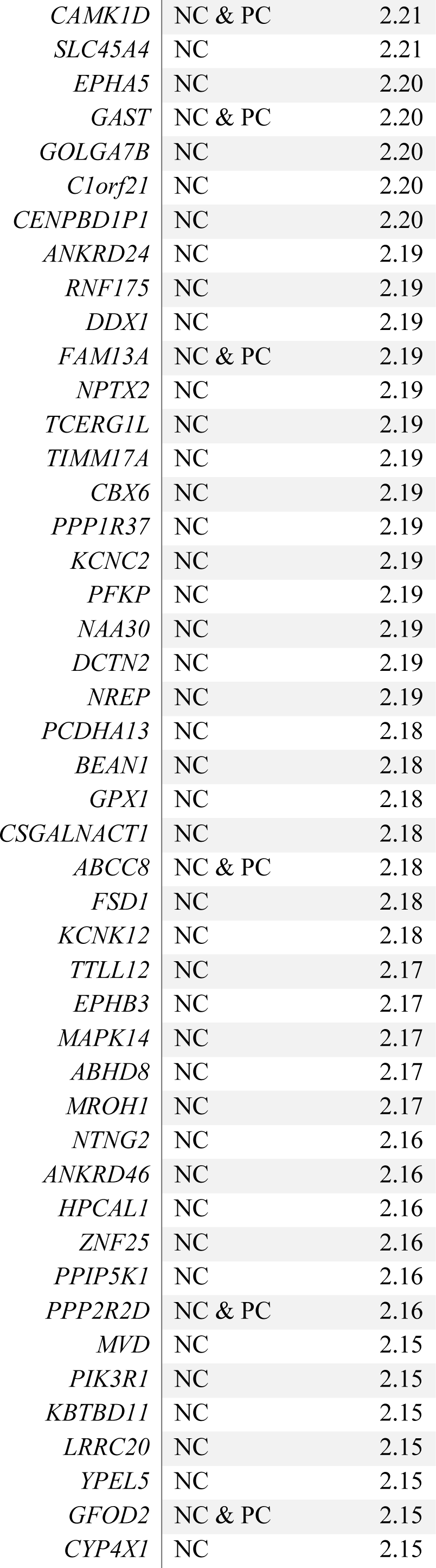
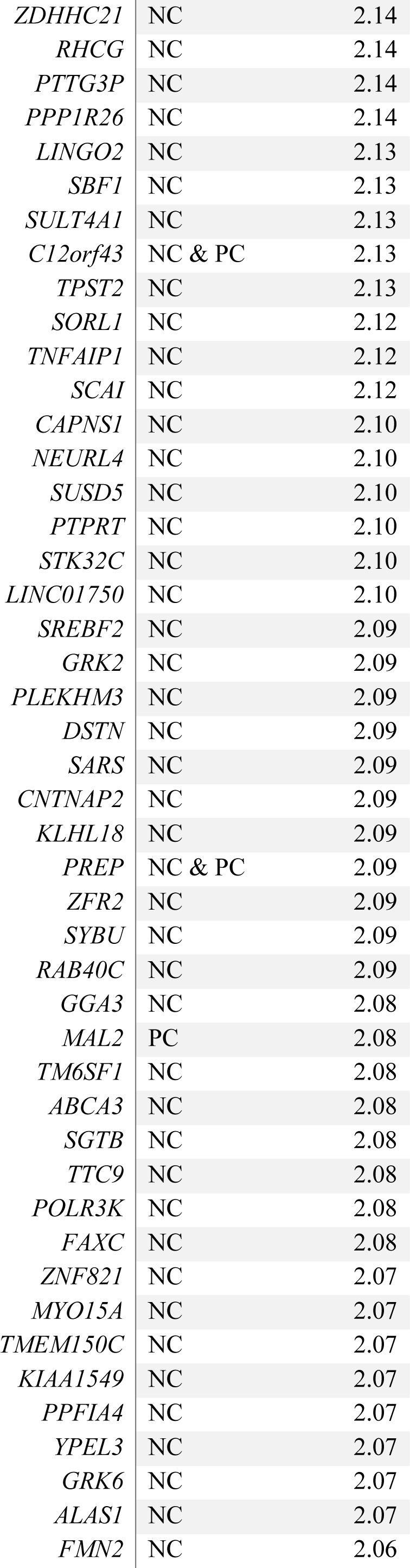
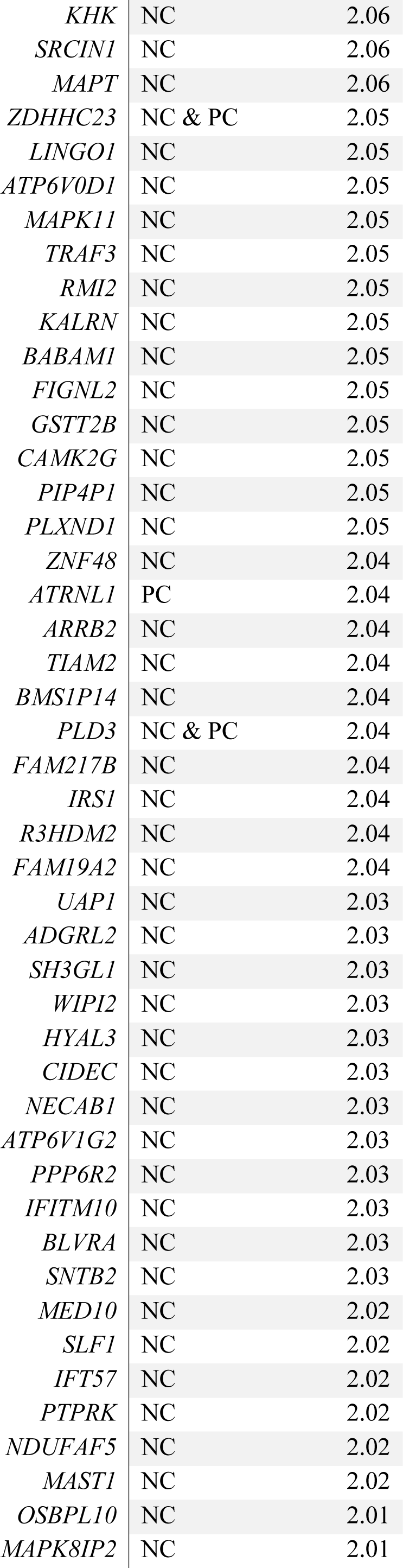
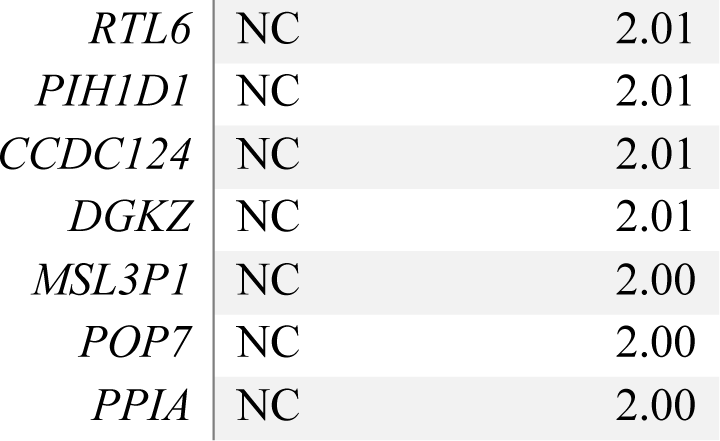
**Negative-correlated genes (blue-cluster) used for enrichment (Cognitive Disability).** Z-score* represents for N genes the Z-score in the probability distribution of the spatial correlation between gene-transcription activity and cognitive disability maps; for connector (C) hubs genes it represents the Z-score in the probability distribution of strength values towards the N-genes cluster.

**Table S9.**
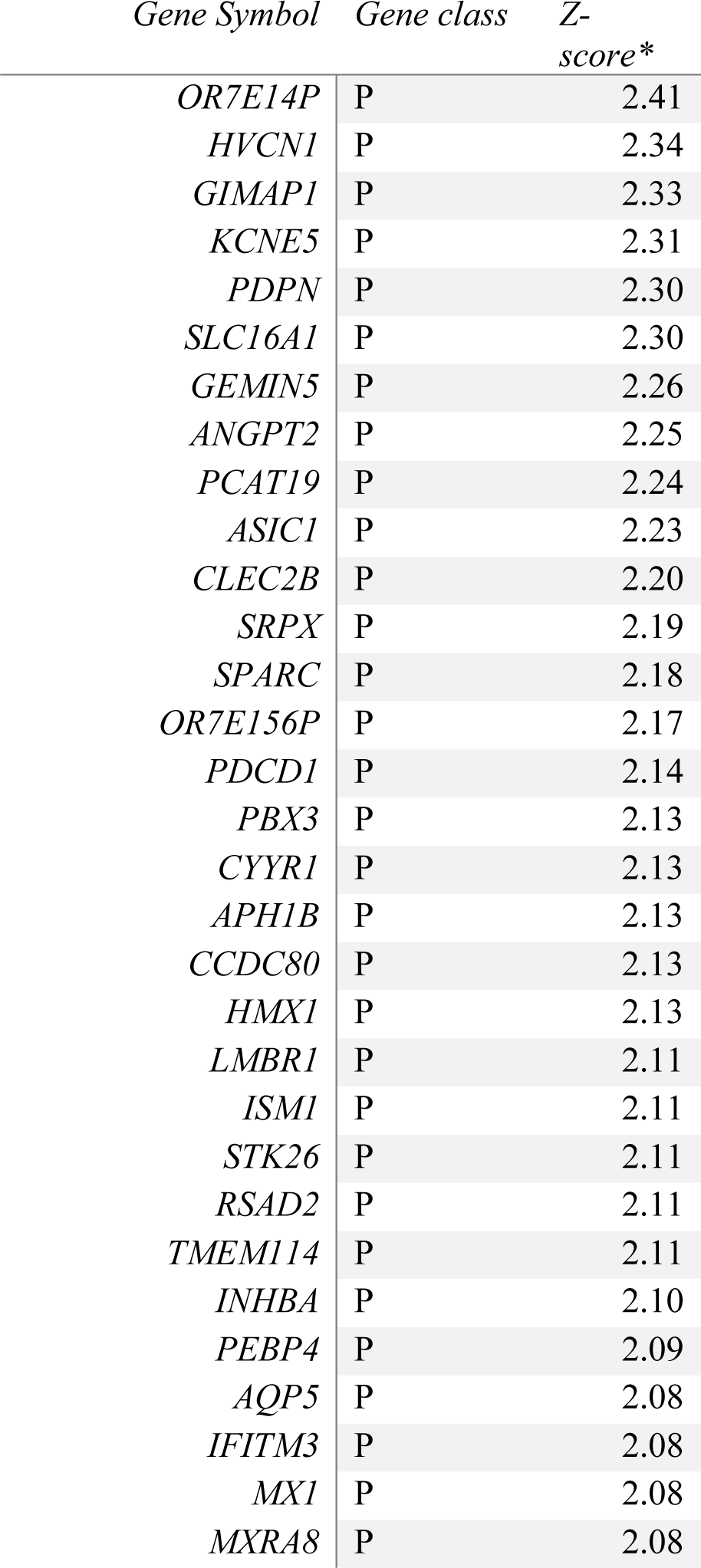
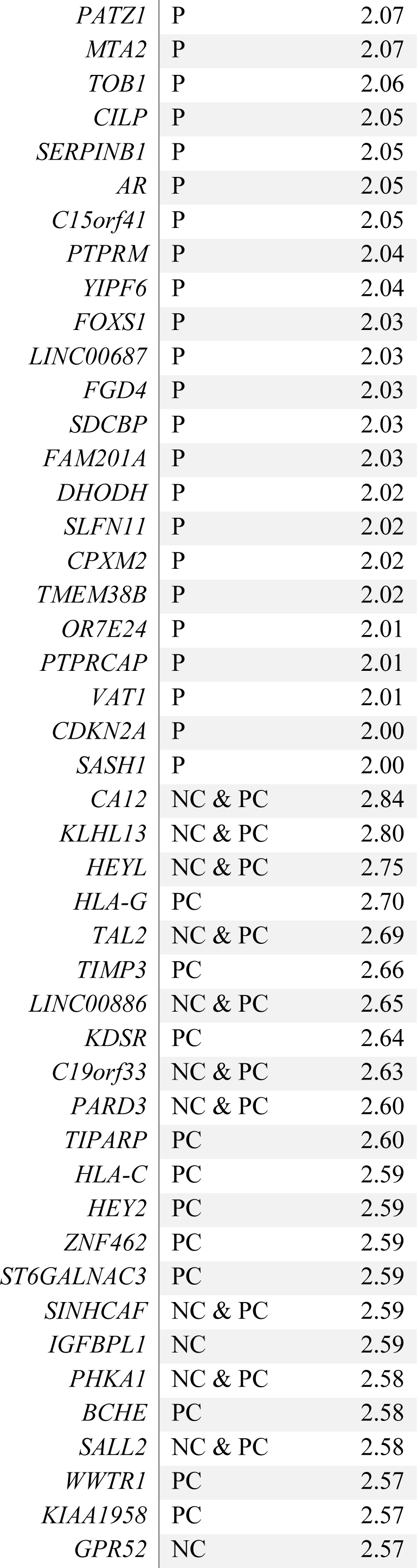
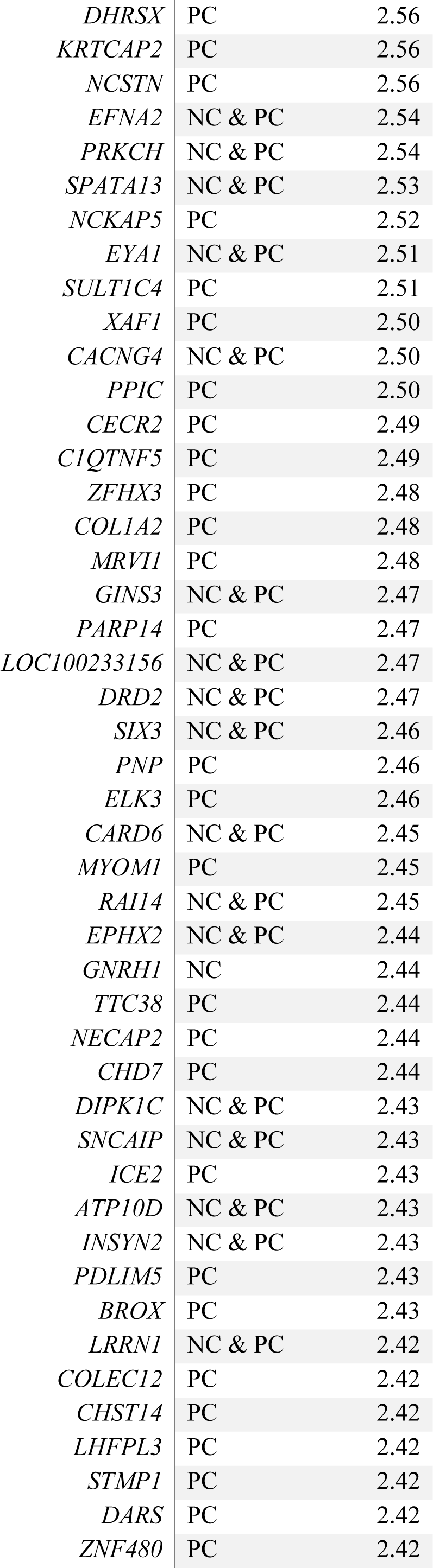
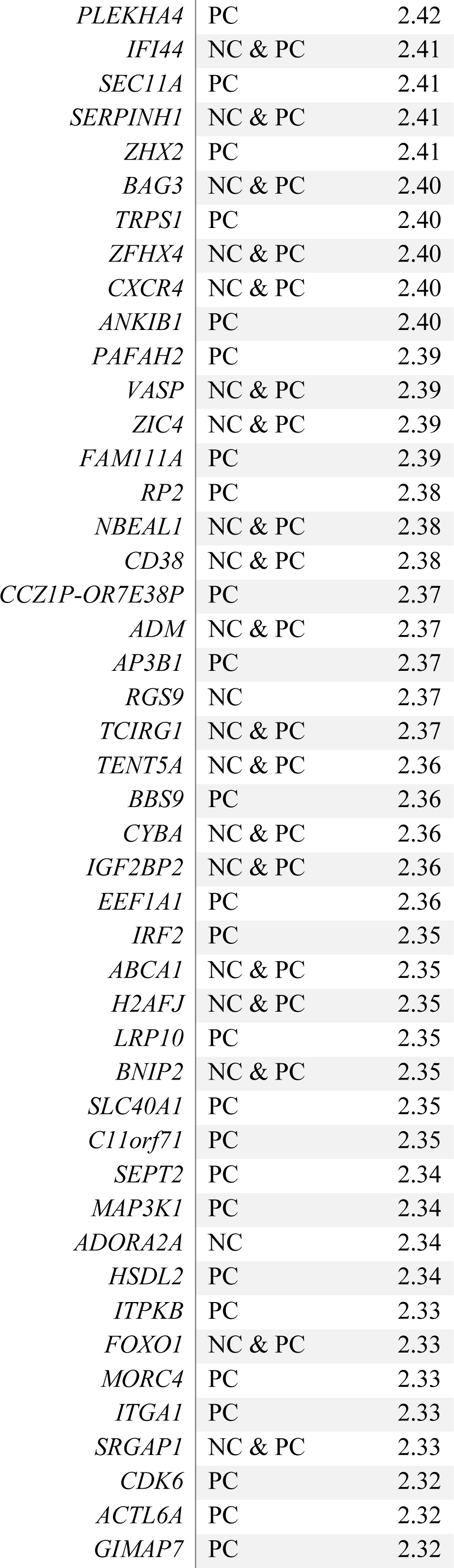
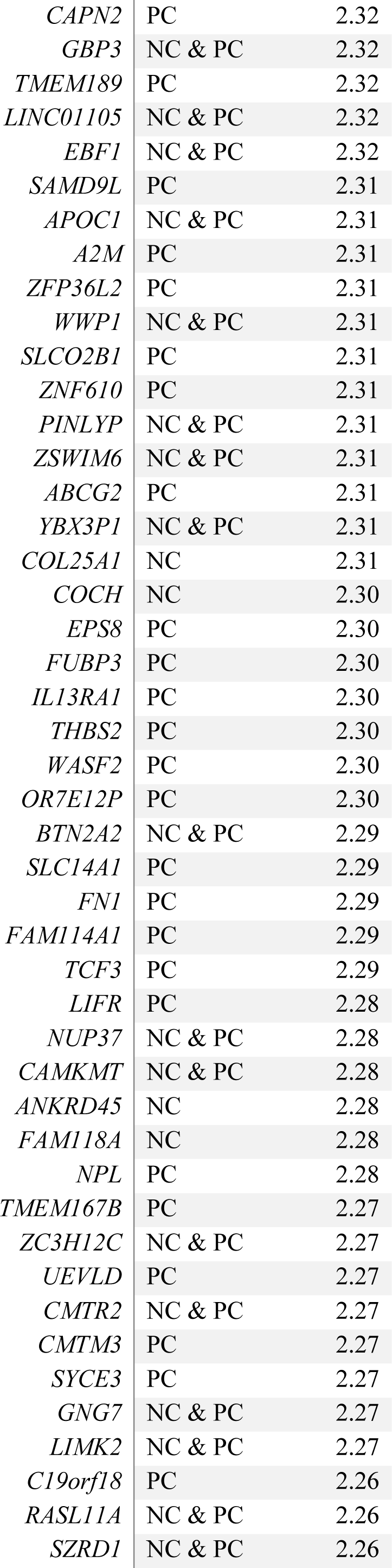
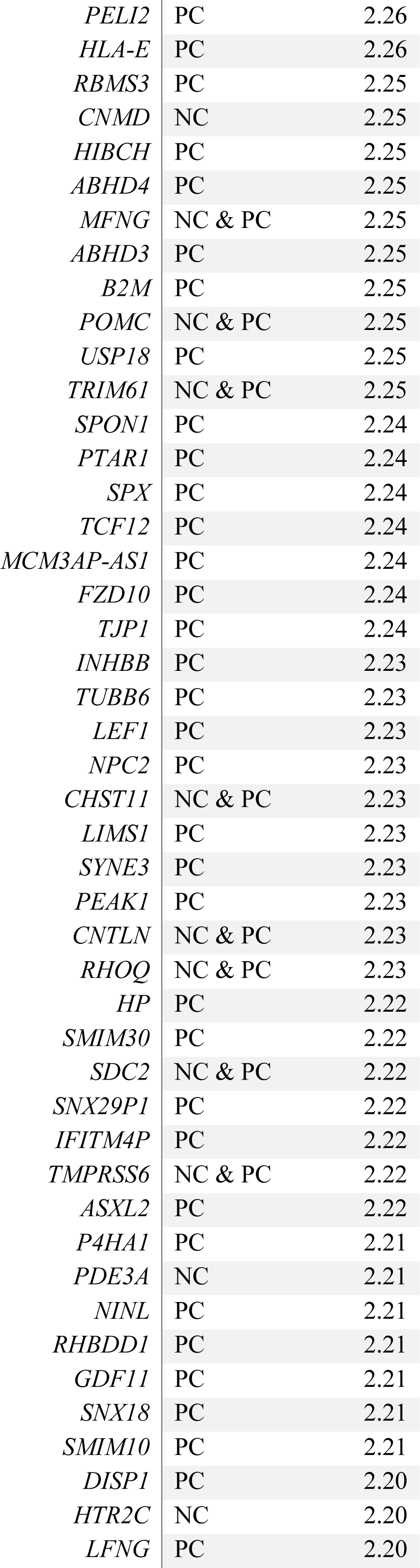
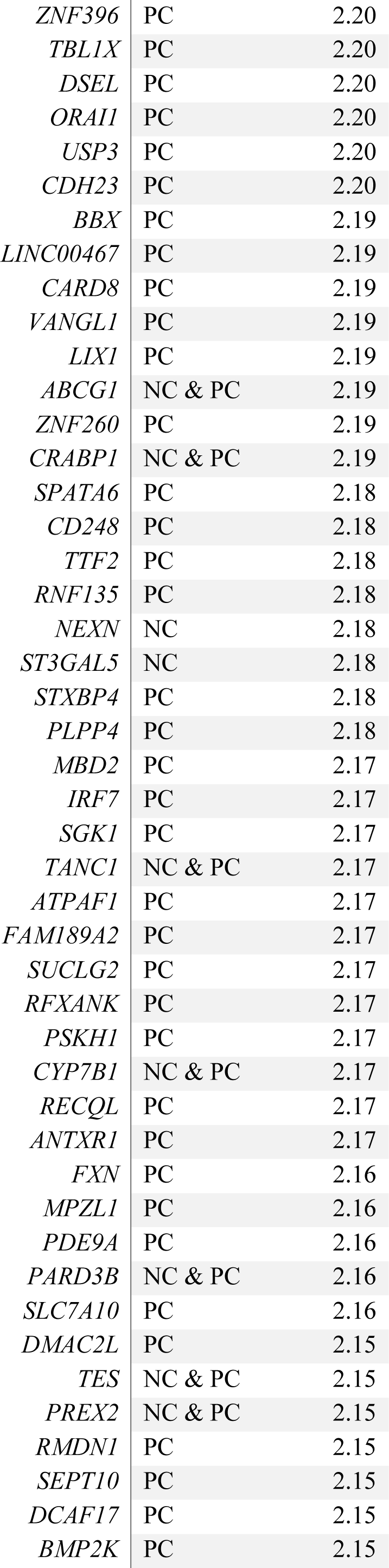
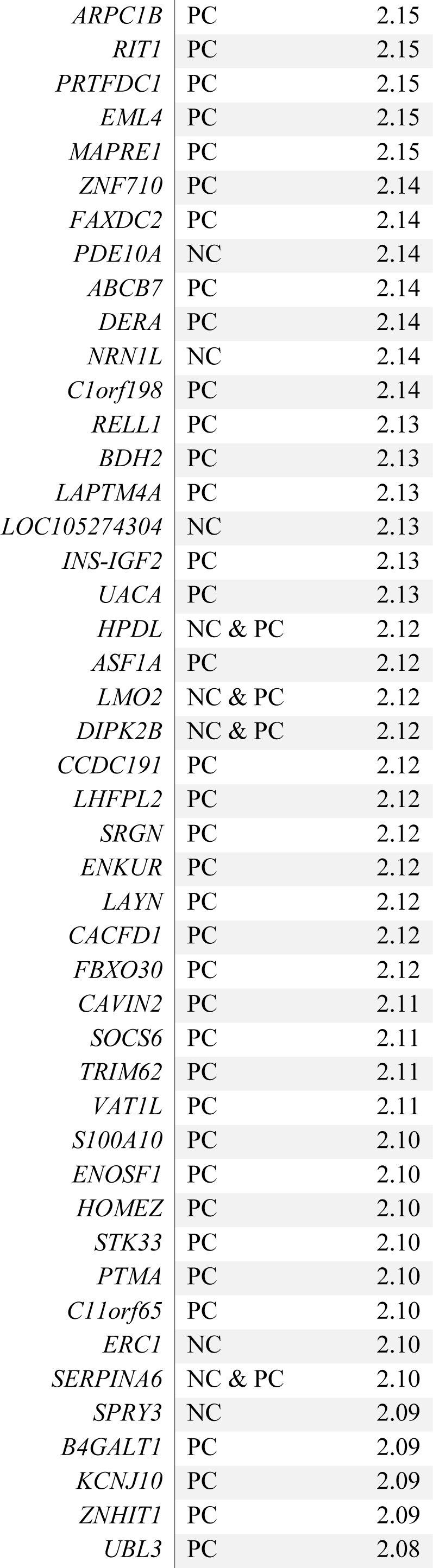
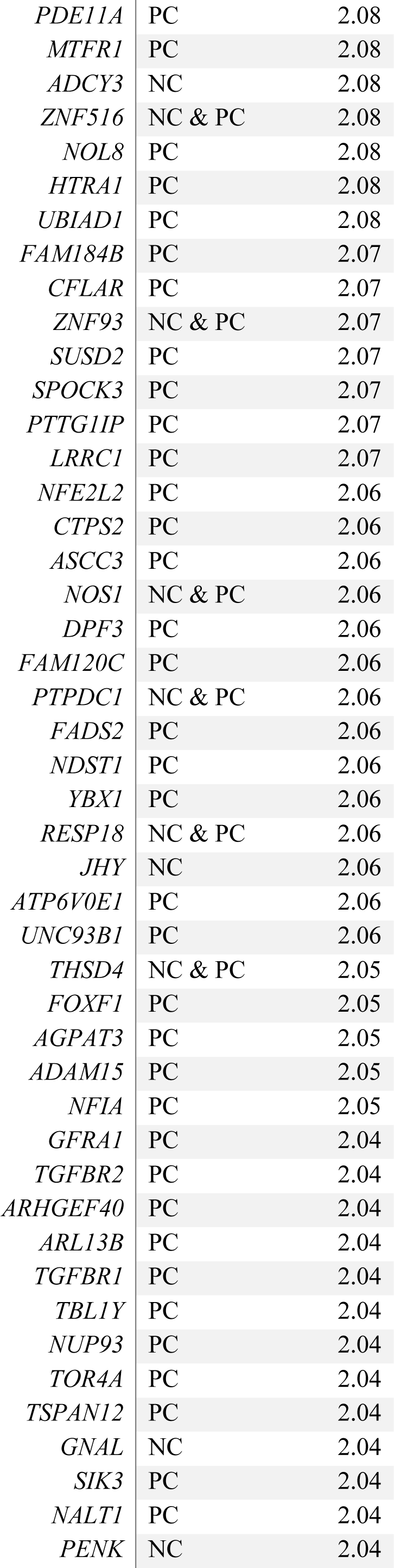
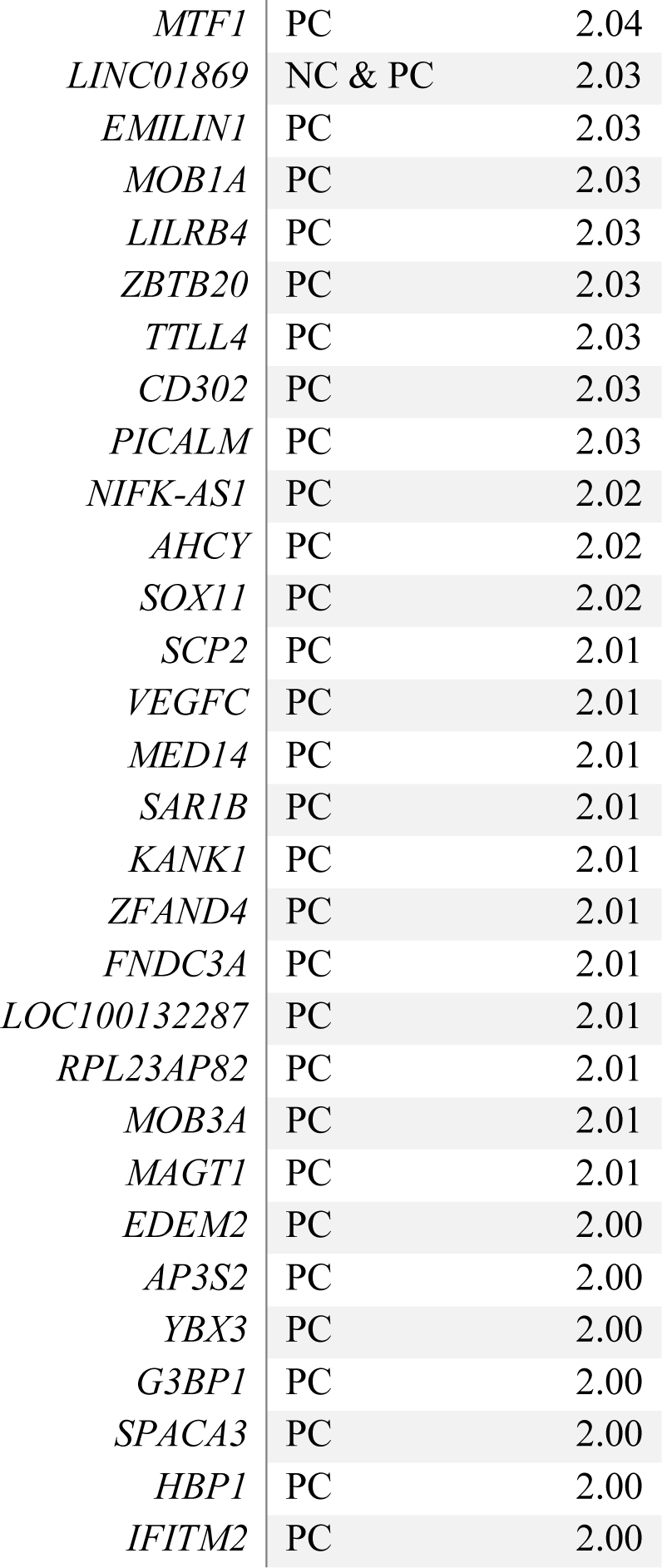
**Positive-correlated genes (red-cluster) used for enrichment (Cognitive Disability).** Z-score* represents for P genes the Z-score in the probability distribution of the spatial correlation between gene-transcription activity and cognitive disability maps; for connector (C) hubs genes it represents the Z-score in the probability distribution of strength values towards the P-genes cluster.

**Table S10.** **Novel genes revealed by the DDS to be implicated in DM1 a.** Three major brain-functions affected in DM1 are highlighted in relation to synaptic vesicle (VES) recycling and dynamics, and the dopamine (DOP) or serotonin (SER) pathways. The statistical significance (p-value) achieved by the surrogate-data is also shown. Because the list of connector hub genes was obtained from the common genes present in the two cohorts, we only represent here one p-value.

| *Gene*<br>*Symbol* | *Function* | *p-value* |  |
| --- | --- | --- | --- |
|  |  | 1<sup>st</sup><br>Cohort | 2<sup>nd</sup><br>Cohort |
| ***Neg-corr genes*** |  |  |  |
| *CALM3* | VES | **0** | **0.0185** |
| *BTBD9* | VES | **0** | **0.0206** |
| *STX1A* | SER, DOP, VES | **0.0005** | **0.0056** |
| *RAB27B* | VES | 0.1129 | **0.0145** |
| *PCLO* | VES | **0.0001** | **0.0113** |
| *BLOC1S6* | VES | **0.0076** | **0.0333** |
| *SNAP25* | SER, DOP, VES | 0.0523 | **0.0082** |
| *DOC2A* | VES | **0.0022** | **0.0120** |
| *ERC2* | VES | **0** | **0.0096** |
| *SYN2* | SER, DOP | **0.0018** | **0.0120** |
| *TSPOAP1* | SER, DOP | **0** | **0.0198** |
| ***Conector-hub genes\**** |  |  |  |
| *DNM1L* | VES | **0.0236** |  |
| *AP2M1* | VES | **0.0007** |  |
| *SH3GL1* | VES | **0** |  |
| *DNM1* | VES | **0** |  |
| *CDK5* | VES | **0.0056** |  |
| *SH3GL2* | VES | **0.0006** |  |
| *SYP* | VES | **0.0003** |  |
| *STXBP1* | VES | **0.0013** |  |
| *RIMS1* | SER, DOP, VES | **0.0003** |  |
| *UNC13A* | VES | **0** |  |
| *RAB3A* | SER, DOP, VES | **0** |  |
| *APBA2* | VES | **0.0003** |  |
| *SLC17A7* | VES | **0.0002** |  |
| *SYT1* | SER, DOP, VES | **0** |  |
| *AMPH* | VES | **0.0002** |  |
| *NRN1* | VES | **0.0001** |  |
| *HTR2A* | VES | **0** |  |
| *GRIN3A* | VES | **0** |  |
| *NAPB* | VES | **0** |  |
| *SYN1* | SER, DOP, VES | **0.0069** |  |

## Footnotes

* AAL regions were eroded with a Gaussian kernel with a full width at half maximum (FWHM) equal to 2 mm, thereby eliminating false-positive sampling sites, i.e.: those that do not belong to the region of interest but to one in the neighborhood.

^†^ To calculate the correlations, the values corresponding to the different regions in the atlas were considered as observations.

